# Macrocyclic Peptide Tools for Huntingtin-bound HAP40

**DOI:** 10.64898/2026.06.02.729325

**Authors:** Esther Wolf, Rebeka Fanti, Tatsuya Ikenoue, Rocher Leung, Renu Chandrasekaran, Matthew G. Alteen, Brandon A. Keith, Suzanne Ackloo, Aled M. Edwards, Derek Wilson, Hiroaki Suga, Rachel J. Harding

## Abstract

Huntington’s disease is a fatal neurodegenerative disorder caused by expansion of a cytosine-adenosine-guanine repeat in exon 1 of the Huntingtin (HTT) gene, resulting in a polyglutamine-expanded HTT protein. Although the genetic cause of Huntington’s is well defined, the molecular functions of HTT and the mechanisms linking polyglutamine expansion to neurodegeneration remain incompletely understood. Huntingtin Associated Protein 40 kDa (HAP40) is a key HTT interaction partner that forms a stable complex with HTT and is increasingly recognized as an important player in the HTT structure-function paradigm. However, investigation of HAP40 biology has been limited by a lack of tools capable of directly targeting endogenous protein. Here, we report the discovery and characterization of a panel of nanomolar-affinity macrocyclic peptides targeting HAP40 identified using Random nonstandard Peptide Integrated discovery platform. We characterized macrocycle binding in vitro using surface plasmon resonance, fluorescence polarization, and hydrogen-deuterium exchange mass spectrometry, revealing selective, high-affinity engagement of distinct epitopes on HAP40. We further demonstrate that these macrocycles engage endogenous HAP40 in cellular lysates, enable selective isolation of HAP40-containing protein complexes using macrocycle precipitations and insights into interaction partners of distinct HTT and HAP40 proteoforms. Together, these macrocycles establish a new toolkit for investigating HTT-HAP40 biology and provide a framework for dissecting HAP40-specific functions relevant to Huntington’s disease pathogenesis.

## Introduction

Huntington’s disease (HD) is a rare neurodegenerative disease occurring in individuals with a *Huntingtin* (*HTT)* allele with more than 35 repeats in its exon 1 cytosine-adenosine-guanine (CAG)-tract.^1^ An inverse relationship exists between CAG-tract length and age of disease onset.^2^ From the time of clinical diagnosis, individuals with HD have a life expectancy of less than 2 decades, and experience debilitating, progressive cognitive, psychiatric, and motor symptoms. Despite the clear genetic association, there are no disease-modifying therapies presently available. Moreover, consensus about *how* the expanded gene and its gene products are causally or mechanistically linked to progressive neurodegeneration and development of HD symptoms is yet to be reached.

In the general population, *HTT* encodes wild-type Huntingtin (HTT), a ubiquitously expressed ∼348 kDa protein with a polyglutamine (polyQ)-tract of ≤36 repeats. Expansion of the CAG-tract in HD results in a polyQ-expanded HTT protein (mHTT). HTT is comprised of Huntingtin-Elongation factor 3-subunit A of protein phosphatase 2A-TOR1 (HEAT) repeats^3^, organizing the protein into two domains: the N-terminal domain (NTD) encompassing the N-HEAT and Bridge regions, and the C-terminal domain (CTD) corresponding to the C-HEAT region.^4^ The precise molecular function of HTT is not clearly defined, but genetic models of HD implicate HTT in various cell pathways, including transcription regulation, proteostasis, cytoskeletal organization, and axonal trafficking.^5–8^ Thousands of interactors are reported for HTT^9,10^, consistent with the role of HEAT repeats mediating protein-protein interactions^11,12^, therefore HTT is hypothesized to be a scaffold protein.^13–15^

A key HTT interaction partner is Huntingtin Associated Protein 40 kDa (HAP40). Cryo-electron microscopy (cryo-EM), crosslinking mass spectrometry (XL-MS), and *in silico* studies of the highly stable monodisperse HTT-HAP40 heterodimer revealed an extensive hydrophobic interface where HTT encases HAP40, while the distal polyQ tract is conformationally flexible and solution-accessible.^16–18^ In contrast, in the absence of HAP40, *in vitro* samples of apo HTT exhibit a flexible conformation, are polydisperse and self-associate, consistent with the absence of a stabilizing binding partner likely present in endogenous settings. A growing body of evidence suggests that HTT and HAP40 co-evolved^19^, their cellular abundances are interdependent^17,20^, and that the dominant protein state of HTT is HAP40-bound.^16,21^ Some studies suggest that HAP40 may influence pathways linked to HTT function, including selective autophagy and vesicular trafficking, potentially through modulation of HTT interactions with factors such as p62 and Rab5.^13,22–24^ Despite these insights, the precise physiological roles of HAP40 and its contributions to HTT-mediated cellular processes or HD pathogenesis remain poorly defined.

Prior studies of HAP40 function have largely relied on HTT-centric approaches, including immunoprecipitation of HTT, to infer associated protein networks. This limits the ability to directly interrogate HAP40-specific interactions and regulatory roles. Some anti-HAP40 monoclonal antibodies (mAbs) have been reported for their use in immunoblotting, however to the best of our knowledge, it is unclear whether they can engage folded, endogenous HAP40. Therefore, tools that enable direct targeting and isolation of HAP40 in its native cellular context are needed to disentangle its physiological function from those of HTT.

We recently reported five HTT-targeting macrocyclic peptides identified by Random nonstandard Peptide Integrated discovery (RaPID) coupled to Flexible in vitro Translation (FIT).^25,26^ These nanomolar affinity macrocycles bound four distinct epitopes across the HTT-HAP40 structure, including sites on HTT and at the interface of HTT and HAP40.^21^ Here we report a panel of nanomolar affinity HAP40-targeting macrocycles. Following identification by RaPID, we characterized in vitro macrocycle binding using surface plasmon resonance (SPR), fluorescence polarization (FP), and hydrogen-deuterium exchange mass spectrometry (HDX-MS). After confirming engagement of endogenous HAP40, we used macrocycle precipitations (MPs) to validate the selectivity of these tools for HAP40 and identify HAP40 interacting proteins. Cross-comparison of MPs with tools targeting different HTT and HAP40 proteoforms identified putative apo HTT-specific and HAP40-specific interactors.

## Results

### Identification of HAP40-targeting Macrocycles

RaPID is a specialized platform that combines RNA-display and flexible in vitro translation (FIT) to enable genetic code reprogramming and incorporation of noncanonical residues, including D-amino acids or chloroacetyl-tyrosine (ClAc-Tyr).^27^ N-terminal ClAc-Tyr and downstream cysteine (Cys) residues can spontaneously react to form stable thioether-bridged cyclized peptides. This generates a library of ∼1 trillion macrocyclic peptides which can be selectively enriched upon binding a target protein, quickly identifying potent and selective binders for targets compared to traditional medicinal chemistry approaches.^28^ Cyclic peptides are excellent chemical tools as they are protease resistant, cheaper than antibodies while retaining similar affinity and selectivity, and may be cell permeable due to their relatively small size.^29^

HAP40 cannot be purified without HTT, so we performed RaPID selections with immobilized, biotin-tagged HTTQ54-HAP40, purified as previously described.^30^ Following five rounds of enrichment, we compared the enriched sequences to those from a prior RaPID screen performed against apo HTT. Sequence analysis of the top-enriched peptides from round five was used to identify binders with potential selectivity for HTT-HAP40 over apo HTT, resulting in eight macrocycles with unique sequences (**Supplementary Table 1**, see **Supplementary Figure 1** for chemical structures).

### Biophysical validation of HAP40-targeting macrocycles reveals nanomolar affinities

Following solid phase synthesis, the eight macrocycles were first assessed by SPR to assess binding for HTT-HAP40 and apo HTT (**Figure 1, Table 1, Supplementary Figure 1**). None of the macrocycles bound apo HTTQ23 with a measurable kinetic affinity at the concentrations tested. Conversely, all macrocycles exhibited nanomolar binding to HTTQ23-HAP40, with K_D_ values ranging from 0.6-47 nM. In particular, macrocycles **2** and **8** had K_D_’s in the single digit nanomolar range of 4.0 and 9.8 nM, respectively. We also observed that macrocycle **6** had a very slow dissociation, nearing the limits of determination, and **5** showed non-specific binding (**Supplementary Figure 2**).

**Figure 1.**
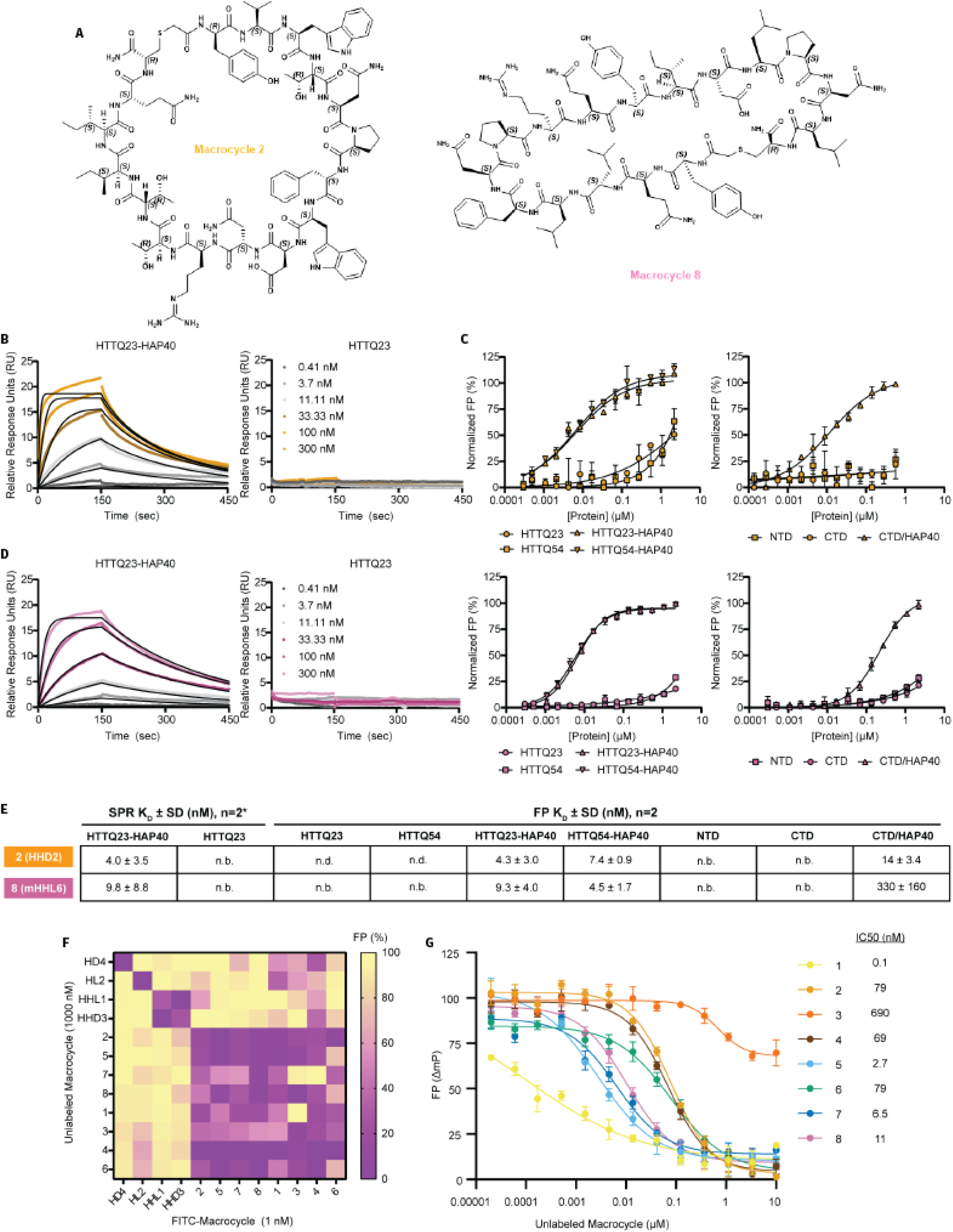
Biophysical Characterization of Macrocycles **2** and **8**. **A**, Structures of macrocycles **2** and **8. B**, SPR with full-length HTTQ23 (n=1)* and HTTQ23-HAP40 (n=2). FP assay with **C**, full-length HTTQ23 or HTTQ54 ± HAP40 (n=2) and **D**, HTT subdomains NTD, CTD, or CTD/HAP40. **E**, summary table of affinity data. K_D_ was not determined (n.d.) when binding curve did not saturate at the concentrations tested. No binding (n.b.) was reported if relative response units (RU) did not exceed 8 RU (∼20% binding) or normalized FP did not reach 50% at concentrations tested. **F**, Single dose displacement matrix with previously reported HTT-HAP40 macrocycles (n=1). **G**, Dose dependent displacement by FP using macrocycle **8** as a tracer (n=1).

**Table 1.** Binding affinities for HTT, HTT-HAP40, and HTT subdomains to different macrocycles determined by SPR and FP.

|  |  | 1 | 2 | 3 | 4 | 5 | 6 | 7 | 8 |
| --- | --- | --- | --- | --- | --- | --- | --- | --- | --- |
| SPR<br>(nM ± SD) | HTTQ23 | n.b. | n.b. | n.b. | n.b. | n.d. | n.b. | n.b. | n.b. |
|  | HTTQ23-HAP40 | 25 ± 20 | 4.0 ± 3.5 | 44 ± 23 | 41 ± 28 | 47 ± 18 | 0.6 ± 0.1 | 29 ± 3.7 | 9.8 ± 8.8 |
| FP<br>(nM ± SD) | HTTQ23 | n.d. | n.d. | n.d. | n.d. | n.d. | n.b. | n.d. | n.b. |
|  | HTTQ54 | n.d. | n.d. | n.d. | n.d. | n.d. | n.b. | n.d. | n.b. |
|  | HTTQ23-HAP40 | n.d. | 4.3 ± 3.0 | 65 ± 10 | n.d. | 13 ± 0.3 | n.d. | 11 ± 4.1 | 9.3 ± 4.0 |
|  | HTTQ54-HAP40 | n.d. | 7.4 ± 0.9 | 32 ± 4.5 | n.d. | 4.4 ± 0.3 | n.d. | 6.9 ± 1.6 | 4.5 ± 1.7 |
|  | NTD | n.d. | n.b. | n.b. | n.d. | n.b. | n.b. | n.d. | n.b. |
|  | CTD | n.d. | n.b. | n.b. | n.d. | n.b. | n.b. | n.d. | n.b. |
|  | CTD/HAP40 | 23 ± 1.5 | 14 ± 3.4 | 94 ± 1.4 | n.d. | 25 ± 0.3 | n.d. | 13 ± 5.9 | 330 ± 160 |

n.d., not determined - binding curve not saturated at concentrations tested
n.b., no detectable binding - normalized FP did not reach 50% at concentrations tested; SPR relative response units did not exceed 8 RU (~20% binding).

Next, using FP, we measured solution-phase binding of fluorescein (FITC)-labeled macrocycles to a suite of HTT proteins, including HTTQ23, HTTQ54, HTTQ23-HAP40, and HTTQ54-HAP40 (**Supplementary Figure 2**). None bound HTT proteins without HAP40. We also observed that **1** (mHHD1), **4** (mHHL3), and **6** (mHHL4) did not have measurable solution-phase affinities at the protein concentrations tested (up to

∼2.5 µM), distinct from our SPR observations, perhaps due to the addition of the FITC label. Macrocycle **3** (mHHL2) had 32 and 65 nM affinity for HTTQ23- and HTTQ54-HAP40, respectively. The remaining macrocycles, including **2** and **8**, had <15 nM affinities for both HTTQ23- and HTTQ54-HAP40 (**Figure 1, Supplementary Figure 2**). Importantly, no macrocycle showed polyQ-dependent binding (polyQ tracts of 23 vs. 54), suggesting the binding mode was independent of this feature of HTT.

We next performed thermal shift assays with our macrocycles and HTTQ23-HAP40. The lowest affinity macrocycles **1, 3, 4**, and **6** yielded minimal thermal stabilization (<1°C) compared to dose-dependent stabilization by **2, 5, 7**, and **8** (**Supplementary Figure 4**).

### Mapping binding interfaces identifies apparent shared binding pocket for all eight macrocycles

To discern where the macrocycles were binding on HTT-HAP40, we explored subdomain binding by FP using our NTD (N-terminal domain), CTD (C-terminal domain), and CTD/HAP40 constructs (**Supplementary Figure 5**).^4^ At the concentrations tested, **4** and **6** did not have a measurable affinity for any of the subdomains, while the remaining macrocycles, including **2** and **8**, engaged CTD/HAP40 with K_D_s in the higher nanomolar range (**Figure 1**). Together, this suggests that **2** and **8** are HAP40 or HTT-HAP40 interface binders. This weakening of affinity could be due to reduced stability of the macrocycle binding interfaces in the context of the subdomain relative to the full-length HTT-HAP40 complex.

In our previous work, we identified five macrocycles with four distinct epitopes on HTT (i.e., HL5) or HTT-HAP40 (i.e., HD4, HL2, HHL1, HHD3). We used these previously described peptides in a single-dose FP displacement matrix to see if any binding sites were shared with the eight new binders. Surprisingly, rather than hitting any of the four previously reported epitopes, we observed that all eight macrocycles displaced one another (**Figure 1**). Since **8** had the largest signal window in previous FP assays (**Supplementary Datasheet 1**), we used it as a displacement assay tracer to corroborate these results in dose-response (**Figure 1**) and observed that all eight macrocycles displaced **8** with IC50 values ranging from 0.1-690 nM. The distinct Hill slopes of the displacement curves suggest different binding modes at a common binding pocket.

### HDX-MS structural characterization localizes binding to HAP40 N-terminal region

To determine where the shared binding site of these eight macrocycles was located, differential hydrogen-deuterium exchange (ΔHDX-MS) was performed across three timepoints using HTT-HAP40 (**Supplementary Figure 8-10**) and the cumulative HDX differences are highlighted in **Figure 2**.

**Figure 2.**
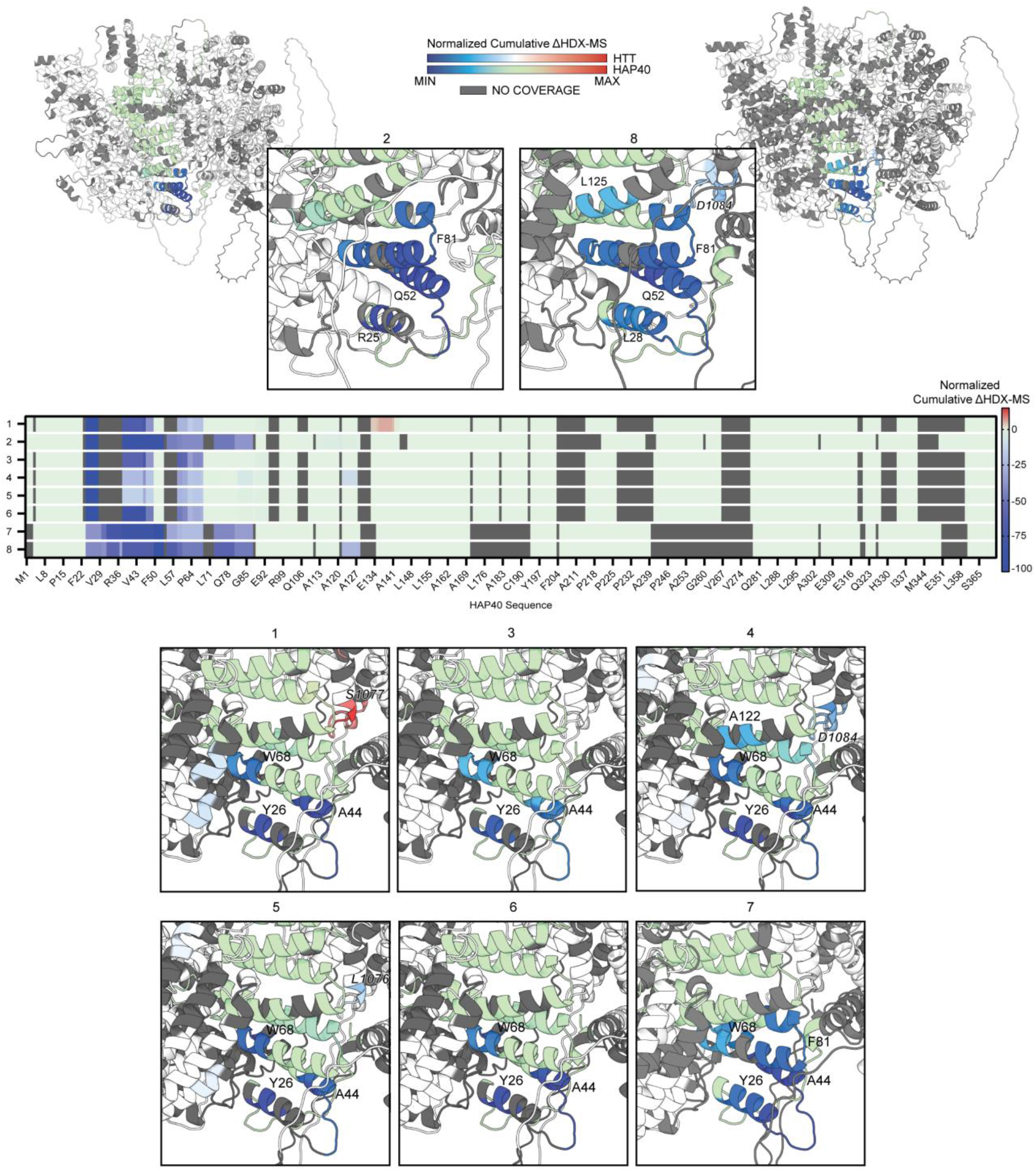
ΔHDX-MS reveals all eight macrocycles target HTT-bound HAP40 N-terminus. **A**, detailed view of macrocycle, ΔHDX-MS footprints **2** and **8. B**, ΔHDX-MS heatmap for linearized HAP40 sequences. **C**, ΔHDX-MS of remaining macrocycles. Differences are reported as cumulative, across three timepoints of labeling (see **Supplementary Figure 9-10** for timepoint-level HDX for HTT and HAP40). To be considered a significant change (increases in red, decreases in blue), it must exceed triple the cumulative propagated error. Insignificant changes are reported in white (HTT) and pale green (HAP40), and missing peptide coverage is grey (see **Supplementary Figure 6-8** for sequence coverage). An AlphaFold3 HTT-HAP40 model was used for visualization.^31^

All eight macrocycles primarily targeted the N-terminal region of HAP40, though with two slightly different footprints across HAP40 and some nearby HTT residues: **2, 7**, and **8**, vs. **1, 3, 4, 5, 6**. Generally, for all macrocycles, HAP40 underwent decreases in HDX at res. 24-28, 38-49, and 59-68, but **2, 7**, and **8** induced more extensive decreases, reaching res. 73-87. The shared HDX footprint for **2, 7**, and **8** points to similar binding mode and may explain why these three macrocycles outperformed others in the biophysical assays (**Figure 1, Supplementary Figure 2-4**).

Uniquely, Macrocycle **1** binding increased deuterium uptake for HAP40 and HTT N-HEAT residues. **1** may have recovered solution-phase affinity for CTD/HAP40 (21 nM) compared to HTT-HAP40 (n.d.) because there was no NTD to undergo destabilization in the truncated construct (**Supplementary Figure 4**). This distinct binding mode, possibly due to its lariat structure (**Supplementary Figure 1**), may factor into the distinct shallow slope observed for **1** in our FP displacement assay (**Figure 1F**). On the other hand, macrocycles **4, 5**, and **8**, induced decreased deuterium uptake at HTT res. 1076-1085 (N-HEAT region), and **4** and **5** induced decreased deuterium uptake at HTT res. 2770-2773 (C-HEAT region). Though impacts to HTT conformational dynamics may be interpreted as orthosteric effects from macrocycles **4** and **5** binding to HAP40, the drop in affinity of **8** for CTD/HAP40 (330 nM) compared to HTT-HAP40 (4.5-9.3 nM) suggests that macrocycle-NTD or NTD-HAP40 contacts may enable tight binding.

Going forward, given the apparent shared binding pocket, we deprioritized macrocycles intractable to FP analysis (1, 3, 4), with trends of nonspecific binding (5), slow dissociation rates (6), and weak thermal stabilization (1, 3, 4, and 6). Of the subdomain FP results, 2 and 8 had the clearest preference for CTD/HAP40 over NTD and apo CTD, and so they became our primary focus.

### Modelling of macrocycle 8 engagement with HAP40

Ideally, to better understand the binding mechanisms of our macrocycles, cryo-EM of the HTT-HAP40 complex would be employed; however, the first ∼80 amino acids of the HAP40 N-terminus are not clearly resolved.^17^ Instead, we sought to model **8** binding to HTTQ23-HAP40 with HADDOCK so that it could be informed by our HDX-MS results.^32,33^ **8** was selected because we had previously observed that linearized macrocycle **8** retained potency (**Supplementary Figure 11**), so, since the unique thioether linkage was not amenable to HADDOCK, linear **8** was used for modeling.

Consistent with the RaPID selection process, where a cognate mRNA would be covalently linked to the C-terminal end of the macrocycle, the modeled C-terminal cysteine of **8** is predicted to be solvent accessible (**Supplementary Figure 11**). The N- and C-termini also remained close together after modeling, suggesting cyclization could be rebuilt, and both the linear and cyclized peptides have similar binding modes. **8** was predicted to form hydrogen bonds with several HAP40 and HTT N-HEAT residues: HAP40 - K40 (via backbone amide), V43 (via backbone amide), A44 (via backbone amide), Q52 (via sidechain carbonyl), and HTT - K1240 (via sidechain amine), and D1243 (via COOH). Though we lacked peptide coverage for those HTT sites in the HDX (**Supplementary Figure 8-10**), HAP40 res. 50-53 underwent the highest decreases in HDX in the macrocycle **8** data set (**Figure 2**), possibly due to the hydrogen bond formed by Q52. This predicted site positions **8** against N-terminal HAP40 residues that face the HTT N-HEAT region, which agrees with the drop-off in FP affinity observed for CTD/HAP40.

Taken together, our data suggests that these 8 macrocycles target the HTT-bound HAP40 N-terminal region with nanomolar affinity, a region that is difficult to visualize using traditional structural techniques but was readily amenable to HDX-MS and *in silico* modeling.

### Validation of HAP40-targeting macrocycles in cellular contexts

To our knowledge, no monoclonal antibody has been robustly validated for immunoprecipitation (IP) of endogenous HAP40. Consistent with this, we tested multiple commercially available antibodies by IP-western blot and did not observe efficient HAP40 pull-down **(Supplementary Figure 13)**, highlighting a current limitation in available tools. Thus, we evaluated the ability of our macrocycles to bind HAP40 in cell lysates **(Figure 3)**. Target engagement was first assessed in HEK293T lysates by macrocycle precipitation (MP) assays using biotinylated analogues of macrocycles **2** and **8**, followed by western blot analysis. Both macrocycles efficiently enriched endogenous HAP40 in the elution, with minimal recovery observed in bead-only controls, confirming that these compounds retain binding in a complex proteomic environment. Further, enrichment of HAP40 was always accompanied by co-precipitation of HTT **(Figure 3A-B)**.

**Figure 3.**
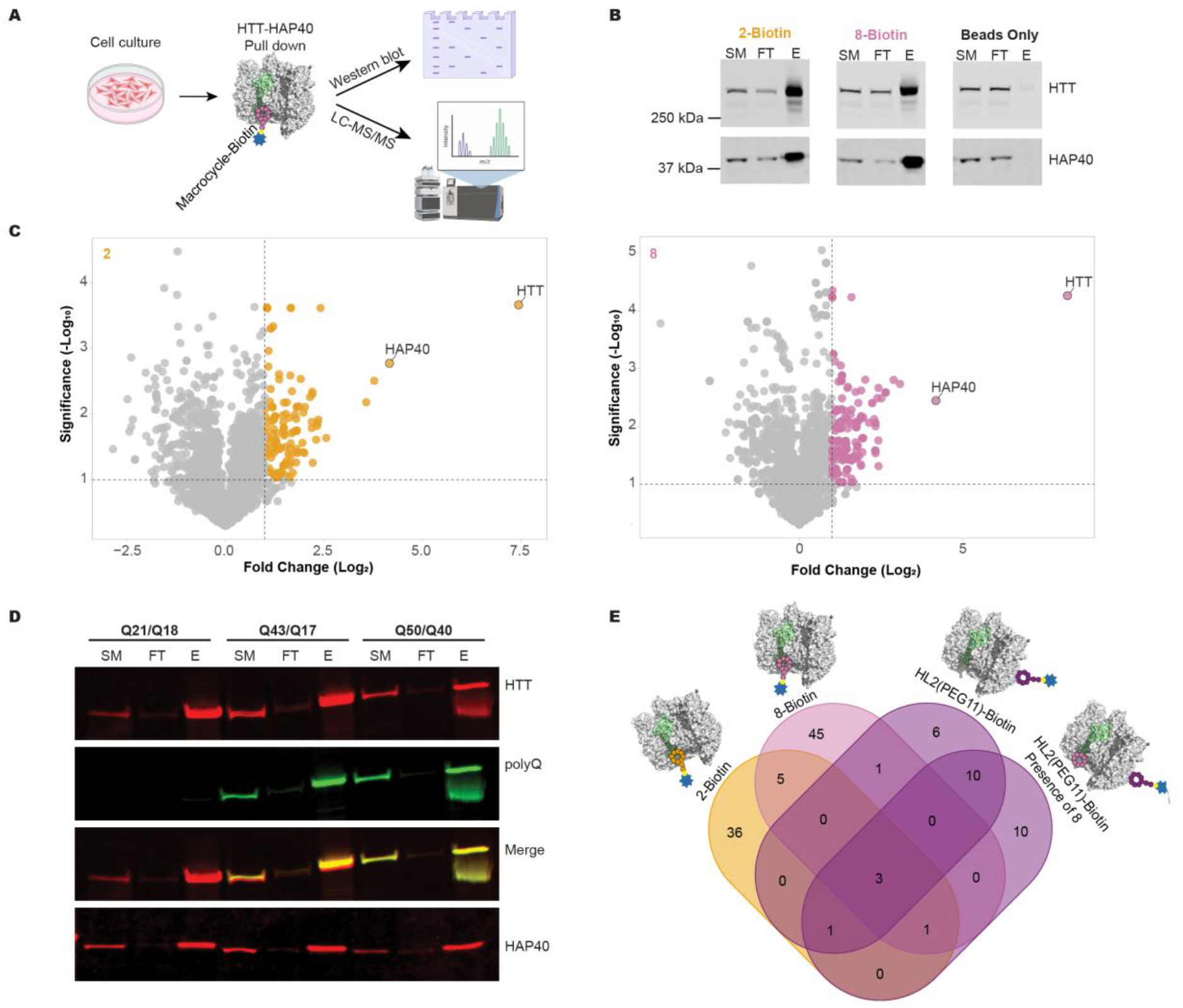
Characterization of macrocycle binding to HAP40 in cells. (A) Schematic of macrocycle pull-down (MP) experiments. Cells were lysed and incubated with biotinylated macrocycles pre-bound to streptavidin beads. Bound proteins were eluted and analyzed by western blot or LC-MS/MS. (B) Western blot validation of HAP40 pulldown. Samples corresponding to starting material (SM) 2%, flow-through (FT) 2%, and elution (E) 100% fractions were analyzed using the anti-HAP40 antibody CH03722 and anti-HTT antibody D7F7, confirming enrichment of HAP40 and HTT in the elution fraction. (C) Volcano plot of LC-MS/MS analysis of macrocycle pulldown (MP) samples, displaying log_2_ fold change (WT vs HTT KO) versus −log_10_ P values derived from normalized protein spectral counts. (n= 3) (D) Macrocycle pulldown performed in fibroblasts harboring different HTT poly-Q lengths, demonstrating enrichment across genotypes. Starting material (SM) 2%, flow-through (FT) 2%, and elution (E) 50%. (E) Venn diagram shows overlap between proteins enriched using HAP40-targeting macrocycles (2-Biotin and 8-Biotin) and the HTT-targeting macrocycle HL2(PEG11)-Biotin, in the presence or absence of macrocycle 8. A ≥3-fold enrichment cutoff relative to HTT knockout controls was applied.

To further evaluate binding selectivity, we extended the MP by coupling to LC-MS/MS-based proteomic analysis. Using HEK293T HTT-null cells^34^, which lack detectable HAP40 spectral counts, as a negative control, this approach identified both HTT and HAP40 as the most enriched proteins for macrocycles **2** and **8**, supporting that these macrocycles selectively target HAP40-associated complexes rather than broadly engaging HTT or nonspecific interactors. Notably, the higher apparent enrichment of HTT relative to HAP40 reflects its large size and correspondingly higher peptide count, which inflates fold-change values when compared to null controls with zero spectral counts, rather than preferential targeting of HTT itself. **(Figure 3C)**.

To determine whether macrocycle-mediated pulldown could be extended to more physiologically relevant systems, we performed MP experiments in HD patient-derived fibroblasts immortalized with human telomerase reverse transcriptase (hTERT; TruHD cells) harboring distinct HTT genotypes (**Figure 3D**). These included control cells with non-pathogenic CAG repeat lengths (Q21/Q18), as well as mutant lines that were heterozygous (Q43/Q17) or homozygous (Q50/Q40) for HD-relevant expansions.^35^ Pulling down HAP40 using biotinylated macrocycle **8**, we observed robust enrichment of HAP40 and HTT in the elution, indicating that macrocycle engagement was maintained and independent on polyQ length, consistent with in vitro observations **(Figure 1D)**.

### Exploring the HAP40 interaction landscape

A previous study identified an N-terminal region of HAP40 as a potential interaction hotspot^36^, which overlaps with our macrocycle binding site. Building on this observation, we asked whether our macrocycles could probe HAP40-associated protein interaction landscapes. We used our previously described HTT-targeting macrocycle, HL2(PEG11)-Biotin, as an affinity reagent for HTT^21^ and pulldowns were performed in the presence or absence of excess HAP40-targeting macrocycle **8**. We reasoned that if macrocycle **8** binds HAP40 at this putative hotspot, its addition would perturb recruitment of proteins dependent on this region. While most proteins remained unchanged, a small subset of proteins appeared to be lost upon addition of macrocycle **8 (Supplementary Figure 14A-B, highlighted in purple)**.

Next, we compared MP-MS datasets generated using HAP40-targeting macrocycles, **2**-Biotin and **8**-Biotin, and the HTT-targeting macrocycle HL2(PEG11)-Biotin, in the presence or absence of **8 (Figure 3E)**. Using a stringent ≥3-fold enrichment cutoff relative to HTT-null controls, we identified candidate HAP40-dependent interactors, defined as proteins enriched in HL2(PEG11)-Biotin pulldowns but absent when HAP40-targeting macrocycles were present. This subset of proteins was also not observed amongst previously reported proteins enriched by apo HTT-selective macrocycle (HL5-Biotin)^21^ **(Supplementary Figure 14C)**, consistent with possible HAP40 association.

Overall, only three proteins were enriched across all conditions: HTT, HAP40, and HEY1. This limited overlap suggests that, aside from the core HTT-HAP40 complex, few proteins are robustly or consistently recovered across macrocycle-based pulldowns. Instead, many HTT-HAP40-associated interactions may be weak, transient, or dependent on epitope and cellular context.

### HTT stabilizes HAP40 but HAP40 does not reciprocally regulate HTT abundance

Given our observation that pull-down of HAP40 consistently co-enriches HTT, together with prior reports showing that knockdown of HTT reduces HAP40 levels^17^, we sought to determine whether this relationship is reciprocal. Importantly, our studies focus on full-length, soluble HTT in a wild-type context rather than HTT aggregation.^37^ To this end, we first examined whether overexpression of HAP40 impacts HTT abundance. In HEK293T cells, ectopic expression of HAP40 did not result in a corresponding increase in HTT levels, indicating that the dependence between these proteins is not bidirectional. Later, treatment with the proteasomal inhibitor MG132 led to a marked accumulation of HAP40, supporting a model in which excess, unbound HAP40 is targeted for proteasomal degradation **(Figure 4A–B)**. Next, we assessed whether reintroduction of HTT could rescue HAP40 levels in an HTT-null background, where HAP40 is only minimally detectable by western blot. Indeed, HTT overexpression restored HAP40 abundance, confirming that HTT stabilizes HAP40 in cells **(Figure 4C)**. This rescue effect was more pronounced in HEK293T cells, whereas comparatively modest to no recovery was observed in U-87MG cells, suggesting potential cell-type-specific differences in HTT-HAP40 regulation.

**Figure 4.**
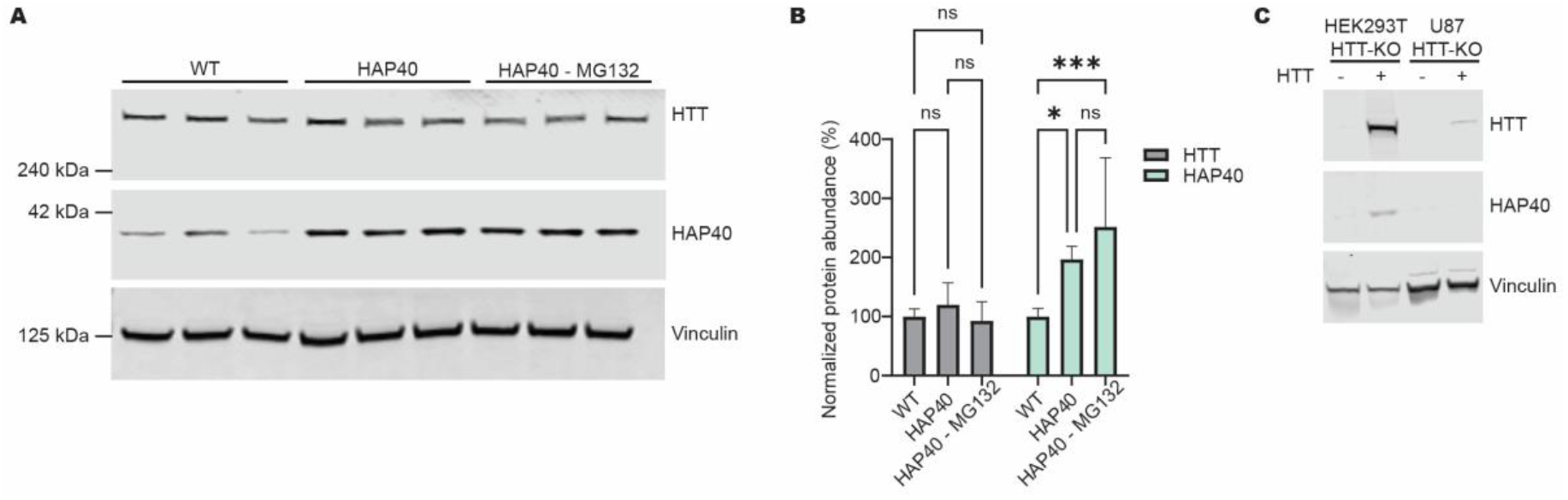
HTT is required for the stabilization and accumulation of HAP40. (A) Representative western blot analysis of WT cells, HAP40-overexpressing cells, and HAP40-overexpressing cells treated with MG132. Vinculin was used as a loading control. (B) Quantification of HTT and HAP40 levels from western blot shown in A, normalized to vinculin and expressed relative to WT. (n= 6) Data are shown as mean ± SD. Statistical significance was determined by ordinary two-way ANOVA followed by Šídák’s multiple-comparisons test. ns, not significant; *p < 0.05; **p < 0.01; *p < 0.001. (C) Western blot analysis of HEK293T HTT-KO and U87 HTT-KO cells with or without ectopic epression of HTT. HAP40 was detected in the HEK293T HTT-KO upon HTT expression, indicating that HAP40 can be rescued in HTT-null cells. Vinculin was used as a loading control (n= 3).

## Discussion

HTT and HAP40 are closely tied, not just in the formation of a high affinity heterodimer, but through their co-evolution and interdependent cellular abundances.^17,19,20^ Indeed, many sources indicate that HAP40 is turned over in the absence of HTT, suggesting that HAP40 has an obligate dependence on HTT heterodimerization.^17,20,36^ Consistent with this, proteomics analysis has shown that the majority of full-length cellular HTT is HAP40-bound.^21^ It is also worth mentioning that co-purification of HAP40 with HTT enabled the cryo-EM studies that solved their atomic resolution structure.^16,17^

Despite this, HAP40 cellular function and how it relates to HTT is not clearly defined beyond stabilization. This is largely because there are few chemical tools available to study it, especially those that are cell permeable and engage endogenous, folded HAP40. To address this gap, we present eight novel, high affinity HAP40-targeting macrocycles that were discovered through the RaPID platform and can be utilized to study HTT and HAP40 function.

Using SPR and FP, we were able to determine these macrocycles bound to HTT-HAP40, in a HTT polyQ-independent manner, and binding was localized to the HTT CTD/HAP40 subdomain complex. Surprisingly, using FP displacement and HDX-MS, we determined that the eight macrocycles impacted a similar region at the N-terminal end of HAP40, though with slightly different HDX footprints. With minimal consensus between the macrocycle amino acid sequences (**Supplementary Figure 11**), and without atomic resolution structures, we could only speculate binding mechanisms or binding mode. To better understand binding mode, we harnessed the utility of HADDOCK to dock macrocycle **8** in an HDX-guided manner. The predicted macrocycle placement was on the N-terminal inner leaflet of HAP40, facing the HTT N-HEAT region.

Using macrocycle precipitations and LC-MS/MS analysis, we observed that macrocycles **2** and **8** were selectively engaging endogenous HTT-HAP40. Also, consistent polyQ-independent binding, macrocycle **8** pulled-down HTT from control and patient-derived fibroblasts.

A recent study by Xu et al. (2022) proposed that the N-terminal helices of HAP40, termed *BΦ* and conserved from Drosophila to humans, may represent a hotspot for mediating protein-protein interactions.^36^ Based on this, we hypothesized that macrocycles targeting the HAP40 N-terminal region could perturb protein-protein interactions associated with this surface. By comparing endogenous proteins pulled down by macrocycles **2** and **8** to our recently published HTT and HTT-HAP40 macrocycles, HL2 and HL5,^21^ we made provisional assignment of candidate interactors to distinct regions of the HTT-HAP40 complex, including HAP40-proximal, apo-HTT-associated, and HL2-pocket-proximal interaction surfaces **(Supplementary Figures 14D-F)**. Although these assignments remain speculative and require validation in future studies using orthogonal approaches, it is encouraging that ARL6IP4, which our analysis predicts to preferentially associate with apo-HTT, was recently reported in a preprint to co-precipitate with HTT from HAP40-knockout cells.^38^ Overall, these analyses highlight the advantage of having complementary affinity reagents that engage distinct sites within the HTT-HAP40 complex. By comparing interaction profiles across macrocycles, we can begin to functionally partition HTT-associated proteins into different classes.

These findings illustrate how chemically defined ligands can be used to functionally partition HTT-HAP40-associated protein networks and probe surface accessibility within native complexes. While the interaction surfaces inferred here were defined in HEK293T cells as a proof of concept, this approach is readily extensible to other cellular contexts, where HTT-HAP40-associated interactions may be cell type-dependent. More broadly, our work highlights the importance of high-quality chemical tools for dissecting protein function, particularly for targets such as HAP40 for which effective biochemical reagents have been lacking.

## Materials and Methods

### Protein Expression and Purification

As described previously^4,30^, HTT variants (Q23 or Q54), HTT–HAP40 complexes, and structure-guided HTT truncations (C-HEAT domain, res. V2095–V3138; N-terminal domain, res. T97–M2069) were expressed in Sf9 insect cells using a baculovirus system. Proteins were FLAG-affinity purified followed by size-exclusion chromatography (SEC). Protein quality control was performed by intact mass spectrometry and SEC post-freeze-thaw from –80 °C. For surface plasmon resonance (SPR) and RaPID assays, biotinylation at the C-terminal HTT Avi-Tag was produced by co-expression with biotin ligase (BirA) in the presence of supplemental biotin.

### Screen for Macrocyclic Peptides that Bind to HTT-HAP40

Macrocyclic peptide selection was performed using the RaPID system, as described previously, with minor modifications. Briefly, mRNA-displayed macrocyclic peptide libraries containing more than 10^12 unique sequences were prepared to identify candidate peptides that bind HTTQ54–HAP40. A puromycin-ligated mRNA library was constructed to encode peptides with N-chloroacetyl-L-Tyrosine (^L^Y-library) or N-chloroacetyl-D-Tyrosine (^D^Y-library) as the initiator amino acid, followed by a random peptide region consisting of 6–15 residues, a cysteine and ending with a short linker peptide. Upon translation of these mRNAs, the chloroacetyl group on N-terminus of the linear peptides spontaneously cyclizes with the downstream cysteine to form thioether-macrocyclic peptides. The cyclic scaffold ensured that the ^L^Y-library and the ^D^Y-library diversified three-dimensional structures. Each cyclic peptide was covalently linked to its corresponding mRNA template via the puromycin linker for later amplification and DNA sequencing. Following four or five rounds of selection, the recovery rate of peptide–mRNA fusion molecules was significantly increased, suggesting that the population of cyclic peptides binding HTTQ54-HAP40 were selectively enriched. Sequence analysis of the respective enriched libraries yielded unique sequences for the peptides. We selected the eight most enriched cyclic peptides for HTTQ54-HAP40 on the basis of the hit frequency for further analysis.

### Chemical Synthesis of Peptides

Macrocyclic peptides were synthesized using standard Fmoc-based solid-phase peptide synthesis (SPPS) on a Syro Wave automated peptide synthesizer (Biotage). The N-terminal Fmoc group was removed with 20% piperidine in DMF, followed by chloroacetylation using chloroacetyl-NHS (ClAc-NHS). The resulting N-ClAc-peptide-Cys-NH-resin (25 μmol scale) was cleaved using a cocktail of 92.5% trifluoroacetic acid (TFA), 2.5% water, 2.5% triisopropylsilane, and 2.5% ethanedithiol to yield the linear N-ClAc-peptide-Cys-NH_2_. After precipitation with diethyl ether, the crude peptide was dissolved in triethylamine containing DMSO and incubated at 25 °C for 1 h to promote macrocyclization. The reaction was quenched with TFA, and the resulting macrocyclic peptide was purified by reverse-phase HPLC (RP-HPLC) using a Shimadzu Prominence system with a linear gradient of solvent A (0.1% TFA in water) and solvent B (0.1% TFA in acetonitrile). Purified peptides were lyophilized and their molecular masses confirmed by MALDI-TOF mass spectrometry (AutoFlex II, Bruker Daltonics).

For biotin and fluorescein labeling, Fmoc-peptide-GSGS-Lys(Mmt)-NH-resin was synthesized by Fmoc SPPS. The Mmt group on Lys was selectively removed using a mixture of 98% dichloromethane, 3% TFA, and 2.5% TIPS. The resulting free amine was conjugated with D-biotin or 5-(iodoacetamido)fluorescein. After N-terminal Fmoc deprotection and chloroacetylation as described above, peptides were cleaved and cyclized by incubation overnight in 5% N,N-diisopropylethylamine in N-methylpyrrolidone at room temperature.

For the Macrocycle 1, which contains Cys(Dpm) for cyclization and Cys(StBu) as a free cysteine, Fmoc-peptide-GSGS-Lys(Mmt)-NH-resin was synthesized similarly. After selective Mmt deprotection, labeling with D-biotin or 5-(iodoacetamido)fluorescein was carried out. Peptides were cleaved, cyclized as above, and the StBu group was removed using tributylphosphine in 10% aqueous solution. Final products were purified by RP-HPLC and lyophilized.

### SPR Assay

SPR was conducted as described previously^21^ using the Biacore 8K SPR system (Cytiva). Briefly, C-terminally biotinylated HTTQ23 or HTTQ23-HAP40 (0.4 mg/mL) was immobilized to a streptavidin (SA) chip to ∼8000 RU (RU_immob_), resulting in an RU_max_ of ∼40-45 RU for the macrocycles. Using Multi Cycle Kinetics (MCK), serially diluted macrocycles were run in biological duplicate in running buffer (10 mM HEPES pH 7.4, 150 mM NaCl, 1 mM EDTA, 0.005% Tween-20 (v/v), 2% DMSO (v/v), 0.02% PEG-3350 (w/v)). K_D_ values were calculated using kinetic fitting by the Biacore Insight Evaluation Software (Cytiva) and were only considered reliable if the concentrations of macrocycle tested spanned the K_D_.

### FP Assays

All FP Assays were conducted using Low Volume 384-well Black Flat Bottom Polystyrene NBS Microplate (Corning Cat 3820), FP buffer (20mM HEPES pH 7.4, 150mM NaCl, 1mM TCEP, 0.005% Tween-20, 1% DMSO, filtered 0.22 µm), and 1 nM FITC-labeled macrocycle. FP was measured using a Synergy Neo2 (BioTek) plate reader set to top optics with ex/em of 485/528 nm. Each biological replicate consisted of a technical triplicate. For affinity determination, HTTQ23, HTTQ54, HTTQ23-HAP40, HTTQ54-HAP40, CTD, NTD, and CTD/HAP40 were prepared in a 2-fold, 14-pt serial dilution with a top protein concentration of 2.8 µM (except for subdomains against macrocycle **2**, 570 nM). Baseline subtracted, normalized curves were fit using Specific Binding with Hill Slope (GraphPad Prism v10.3.0). For the macrocycle matrix FP assay, a final concentration of 1 µM of each unlabeled macrocycle was mixed with 10 nM HTTQ23-HAP40 in technical duplicate. The data was baseline subtracted and normalized. For the macrocycle **8** displacement assay, unlabeled macrocycles were prepared in a 14 pt, 3-fold serial dilution starting from 10 µM and mixed with a final concentration of 10 nM HTTQ54-HAP40. Baseline subtracted curves were fit using [Inhibitor] vs. response -- Variable slope (four parameters) to determine the IC50 values of each unlabeled peptide (GraphPad Prism v10.3.0).

### HDX-MS

HDX-MS was performed as previously described.^21^ Briefly, 12 µM HTTQ23-HAP40 ± macrocycle 1, 2, 3, 4, 5, 6 and HTTQ54-HAP40 ± macrocycle 7, 8 were prepared in a 1:2 ratio in storage buffer (20 mM HEPES pH 7.0, 300 mM NaCl, 2.5% glycerol (v/v), 1 mM TCEP, 2% DMSO (v/v)) and held at room temperature for the duration of the experiment. Samples were labeled for 0, 0.25, 1, and 10 mins using deuterium buffer (10 mM Potassium Phosphate, pD 7.5) at 20 °C (except macrocycle 2 samples which were labeled for 0, 0.5, 10, and 30 mins). Then, samples were quenched in harsh conditions (0.4 M TCEP, 6 M Gdn-HCl, 100 mM Potassium Phosphate pH 2.5, filtered 0.22 µm) for 2 mins at 0°C followed by mixing 1:1 with diluent (100 mM Potassium Phosphate, pH 2.5). Samples underwent on-line digestion at 15 °C using a Nep2-Pep enzyme column (Affipro) and then peptides were desalted (ACQUITY UPLC BEH C18 VanGuard Pre-column, Waters) and separated (ACQUITY UPLC BEH C18 Column, Waters) prior to MS analysis. Peptide identification and HDX data processing were performed using ProteinLynx Global Server (PLGS) and DynamX, respectively. Data was further processed using Microsoft Excel and visualized using PyMOL and GraphPad. An AlphaFold3 model of HTTQ23-HAP40 was used to ullustrate the cumulative ΔHDX-MS heatmaps. Cumulative differences were considered significant if they exceeded triple the cumulative propagated error.

### HADDOCK Modeling

Macrocycle **8** was fabricated and sculpted in PyMOL v3.1.4.1 (Schordinger, LLC). Also using PyMOL, the sequence of HTTQ23-HAP40 (AlphaFold 3) was concatenated (3515 aa.). The HTTQ23-HAP40 complex and macrocycle were submitted to the HADDOCK v2.4 Web Server (EASY mode) as two molecules, using residues which underwent significant changes in HDX as inputs for Active Residues. Of the 96 structures grouped into 10 clusters, **Supplementary Table 2** highlights the top 5 clusters, with cluster 3 scoring the most favorably (**Supplementary Figure 12**). HADDOCK score is a weighted sum of Energies (Van der Waals, intermolecular electrostatic energy, desolvation, and ambiguous interaction restraints). Cluster 3 was prioritized for this manuscript.

### Thermal Shift Assay

All 8 macrocycles were tested at a 5 pt, 2-fold serial dilution, with a top concentration of 10 µM against 0.2 mg/mL HTTQ23-HAP40 in 20 mM HEPES pH 7.4, 150 mM NaCl, 0.005% Tween-20, and 2% DMSO. The samples were loaded into a low volume, 384-well PCR microplate (Axygen® REF# PCR-384-LC480-W-NF, Corning) with a final concentration of 5X SYPRO Orange (CAT# S6650, Thermo Fisher Scientific). The assay was run using a LightCycler® 480 Instrument II with a thermal ramp of 20-95 °C (4 °C/min) and fluorescence detection of 580 nM. The experiment was performed in biological duplicate, reported as the average thermal shift (ΔTm ± SD).

### Cell culture

HTERT-immortalized control (Q21/Q18) and HD patient–derived fibroblasts (Q43/Q17 and Q50/Q40) were cultured in Dulbecco’s modified Eagle’s medium (DMEM; Life Technologies, #10370) with 15% (v/v) FBS (Gibco, #12484-028), 1× GlutaMAX (Life Technologies, #35050), and 1× penicillin-streptomycin (PenStrep) antibiotics. U-87MG wildtype and HTT knockout, HEK293T wildtype and HTT null were cultured in Dulbecco’s modified Eagle’s medium (DMEM; Life Technologies, #10370) with 10% (v/v) FBS (Gibco, #12484-028) and 1× penicillin-streptomycin (PenStrep) antibiotics.The cells were grown at 37°C with 5% CO2.

### Biotinylated Macrocycle Pull-Down

Biotinylated macrocycle-pull down has been previously described.^21^ Briefly, Macrocycles were pre-conjugated to streptavidin beads (Cytiva, 17-5113-01) by incubation with a 30 μL bead slurry per macrocycle precipitation (MP) in lysis buffer (10 mM Tris-HCl, pH 7.9, 150 mM NaCl, 0.1% NP-40, 1× protease inhibitor cocktail (Roche, 11873580001), 1:1,000 benzonase nuclease, and 2 mM MgCl_2_) for 1 h at 4 °C, using a final macrocycle concentration of 31.25 μM. Beads were washed three times with lysis buffer to remove unbound macrocycles. Cells were washed twice with PBS, harvested by centrifugation (1,500 rpm, 5 min), and snap-frozen. Pellets, each derived from a confluent 150 mm cell culture dish, were resuspended in 1 mL of lysis buffer and incubated at 4 °C for 30 minutes with end-over-end rotation. Lysates were clarified by centrifugation (20,000 × g, 15 min, 4 °C), and protein concentration was determined by BCA assay (Thermo Fisher Scientific, 23225). Macrocycle-conjugated beads were incubated with 5 mg of lysate (500 μL at 10 mg/mL) for 1 h at 4 °C with rotation. Beads were collected by centrifugation (3,000 rpm), and the flow-through was retained. Beads were washed once with 1 mL Wash Buffer 1 (10 mM Tris-HCl, pH 7.9, 100 mM NaCl, 0.1% NP-40) and twice with Wash Buffer 2 (10 mM Tris-HCl, pH 7.9, 100 mM NaCl). Bound proteins were eluted with 200 μL 0.5 M NH_4_OH and flash-frozen in liquid nitrogen.

### Preparation and Analysis of MP Samples for LC-MS/MS

Sample preparation and LC–MS/MS analysis were performed as previously described^21^ at the SPARC BioCentre (The Hospital for Sick Children, Toronto, ON, Canada). Samples were reduced with DTT (10 mM, 60 °C, 1 h), alkylated with iodoacetamide (20 mM, room temperature, 45 min, dark), and digested overnight at 37 °C with trypsin (Pierce). Peptides were dried by vacuum centrifugation, desalted using C18 ZipTips (Millipore) on a DigestPro MSi (Intavis), dried again, and resuspended in Buffer A (0.1% formic acid in water). Peptides were analyzed on an EASY-nanoLC 1200 system coupled to an Orbitrap Fusion Lumos Tribrid mass spectrometer (Thermo Fisher Scientific). Separation was performed on a 75 μm × 50 cm PepMap RSLC EASY-Spray column (2 μm C18 beads) with a 75 μm × 2 cm Acclaim PepMap 100 pre-column at 250 nL/min, using a gradient of 3–20% Buffer B (0.1% formic acid in acetone) over 18 min, 20– 35% over 31 min, followed by a 2 min ramp to 100% and a 9 min hold. MS1 spectra were acquired at 120,000 resolution (m/z 375–1,500; AGC 4 × 10^5^; IT 50 ms), with data-dependent MS2 acquired in the ion trap (AGC 1 × 10^4^; IT 10 ms; isolation window 0.7 m/z; HCD NCE 30; dynamic exclusion 10 s). Raw data were processed using PEAKS Studio (Bioinformatics Solutions Inc.) and Proteome Discoverer (v3.1.1.93), searching against the human UniProt database (UP000005640, July 2024). Parent and fragment mass tolerances were set to 50 ppm and 0.02 Da, respectively, with fully tryptic specificity and up to three missed cleavages allowed. Normalized spectral counts were extracted using Scaffold DDA software, and data visualization was performed using VolcaNoseR.^39^

### Immunoblotting

Immunoblotting was performed as previously described.^21^ In short, cells were lysed in RIPA buffer (Thermo Fisher, 89900) supplemented with 1X protease inhibitors (Roche, 11873580001) for 30 min on ice. Lysates were clarified by centrifugation (20,000 × g, 15 min, 4 °C), and protein concentration was quantified prior to loading. 20–30 µg were resolved on 4–12% Bis-Tris gels (Invitrogen) in MOPS running buffer (1 M MOPS, 1 M Tris base, 69.3 mM SDS, 20.5 mM EDTA) at 150 V for 90 min and transferred to 0.2 µm PVDF membranes using transfer buffer (25 mM Tris base, 192 mM Glycine, 20% Methanol) at 45 V for 2 h. Membranes were blocked in 5% nonfat milk in PBS-T (PBS + 0.05% Tween-20) for 1 h and incubated overnight at 4 °C with primary antibodies diluted in blocking buffer (5% BSA, 0.02% NaN_3_): anti-HTT (1:1,000; 1HU-4C8 or D7F7), anti-HAP40 (1:500; CH03722), anti-Vinculin (1:2,000; Santa Cruz sc-73614). Following washing (3 × 5 min in PBS-T), membranes were incubated with IRDye secondary antibodies (LI-COR; 1:5,000) for 1 h at room temperature, washed again, and imaged using a LI-COR Odyssey CLx scanner.

### Immunoprecipitation

Immunoprecipitations were performed as previously described.^40^ Cells were lysed in IP lysis buffer (Thermo Fisher Scientific, 87788) supplemented with 1× protease inhibitor cocktail (Roche, 11873580001). Soluble material was cleared by centrifugation at 20,000 x g for 15 min at 4°C. Clarified lysates were adjusted to 1.0 mg/mL with lysis buffer and incubated with 2 µg of antibody pre-coupled to 30 µL protein A/G magnetic beads (Thermo Fisher Scientific, 10001D, Thermo Fisher Scientific, 10004D) for 1 h at 4°C with end-over-end rotation. Starting materia and flowthrough fractions were retained for subsequent analysis. Immunoprecipitated proteins were eluted by boiling the beads in SDS-PAGE loading buffer for 5 min at 95°C, resolved by SDS-PAGE, and analyzed by immunoblotting.

### Allele Separation Immunoblotting

Allele-separating immunoblotting was performed as previously described.^41^ Lysates (15–35 µg total protein) were resolved on 3–8% Tris-acetate gradient gels (Invitrogen, EA03785BOX) at 120 V for 24 h at 4 °C, until the 250 kDa molecular weight marker migrated out of the gel. Tri-acetate running buffer (25 mM Tris base, 190 mM glycine, 3.5 mM SDS, and 10.7 mM β-mercaptoethanol) was pre-chilled and replaced twice during the electrophoresis to maintain optimal separation. Following electrophoresis, proteins were transferred onto 0.2 µm PVDF membranes (Bio-Rad) using transfer buffer as described above at 45 V for 2 h. Membranes were blocked for 1 h at room temperature in 5% nonfat milk prepared in PBS-T. Membranes were incubated with primary antibody against anti-polyQ MW1 (1 µg/mL; DSHB) and anti-HTT EPR5526 (1:1000; Abcam; ab109115), diluted in blocking buffer. Secondary antibody incubation and fluorescence detection were performed as described above.

### HAP40 rescue experiment

HEK293T HTT-null cells and U-87 HTT-knockout cells were transfected with 1 μg of pBacMam2-DiEx-LIC-C-flag_huntingtin_full-length_Q23 plasmid (Addgene Plasmid, #111726) using X-tremeGENE DNA Transfection Reagent (Roche, 6366244001), according to the manufacturer’s instructions. Cells were incubated for 48 h post-transfection, then lysed and analyzed by western blotting as described above.

### HAP40 overexpression and MG132 treatment

HEK293T cells were seeded in 6-well plates and transfected with 1 μg of HAP40_pcDNA3.1 plasmid using X-tremeGENE DNA Transfection Reagent (Roche, 6366244001), according to the manufacturer’s instructions. Cells were incubated for 24 h post-transfection. After 24 h, one set of HAP40-overexpressing cells was collected, while a second set was treated with 10 μM MG132 for 6 h before collection. Wild-type, HAP40-overexpressing, and HAP40-overexpressing + MG132-treated cells were lysed and analyzed by western blotting as described above.

## Acknowledgements

The FP7 WeNMR (project# 261572), H2020 West-Life (project# 675858), the EOSC-hub (project# 777536) and the EGI-ACE (project# 101017567) European e-Infrastructure projects are acknowledged for the use of their web portals, which make use of the EGI infrastructure with the dedicated support of CESNET-MCC, INFN-LNL-2, NCG-INGRID-PT, TW-NCHC, IFCA-LCG2, UA-BITP, TR-FC1-ULAKBIM, CSTCLOUD-EGI, IN2P3-CPPM, SURFsara and NIKHEF, and the additional support of the national GRID Initiatives of Belgium, France, Italy, Germany, the Netherlands, Poland, Portugal, Spain, UK, Taiwan and the US Open Science Grid. We thank M. E. MacDonald and I. S. Seong from Massachusetts General Hospital for their gift of HTT null HEK293T cells. We thank R. Truant from McMaster University for the gift of TruHD patient-derived fibroblasts. We thank Craig Simpson at SPARC BioCentre and Greg Wasney at the Structural & Biophysical Core Facility at the Hospital for Sick Children, Toronto, Canada for their assistance with mass spectrometry and biophysical instrumentation. Finally, we thank CHDI Management / CHDI Foundation for providing the anti-HAP40 mAb CHDI-90004692 used in Western blot analyses.

## Author Contributions

Conceptualization, R.J.H., H.S., and M.G.A.; Data curation: E.W., R.F., T.I., R.L., M.G.A., B.A.K., and R.C.; Formal analysis: E.W., R.F., T.I., and R.L.; Funding acquisition: R.J.H, H.S., D.W., and A.M.E.; Investigation and Methodology: E.W., R.F., T.I., R.L., and R.C.; Project administration and Supervision: R.J.H, H.S., D.W., and A.M.E.; Resources and Software: R.C., T.I., D.W.; Writing – original draft: E.W., R.F., and R.J.H.; Writing – review & editing: E.W., R.F., T.I., R.L., S.A., A.M.E, and R.J.H.

## Data Availability Statement

The FP and proteomics data supporting this article have been included as part of the Supplementary Information. All HDX-MS files are available on Zenodo at 10.5281/zenodo.20173864, 10.5281/zenodo.20336751, 10.5281/zenodo.20337207. All LC-MS files are available on MassIVE at MSV000101852 (https://massive.ucsd.edu/ProteoSAFe/dataset.jsp?task=5db2aa9bf96d4eb6abd1f2e4ea295682).

