## Supplemental Data for "Macrocyclic Peptide Tools for Huntingtin-bound HAP40"

### Supplementary Tables and Figures

**Supplementary Table 1.** Macrocycle Nomenclature and Sequences

| Number | Linear Sequence* |
|---|---|
| 1 | $^{D}$YLCHYFTNPYEYWLSCC |
| 2 | $^{D}$YVWTNPFWDNRTTIIQC |
| 3 | YRIWFNPFHQGWTVVVC |
| 4 | YAYGDRHLFRHLFDDRC |
| 5 | $^{D}$YIRWTNPPFGRYYRWSC |
| 6 | YGDRNLDLLLRIRNRFC |
| 7 | YTNWYWLHNRFSTYVDC |
| 8 | YQLLFNPRQYIDLPNLC |

*All macrocycles are cyclized through a thioether between the N-terminus and C-terminal Cys sidechain.
$^{D}$N-terminal Tyr is D stereoisomer, rather than L.

**Supplementary Table 2**. HADDOCK clustering results

| | CLUSTER 3 | CLUSTER 9 | CLUSTER 10 | CLUSTER 2 | CLUSTER 8 |
|---|---|---|---|---|---|
| **HADDOCK score** | -36.6 +/- 6.6 | -24.5 +/- 3.8 | -22.9 +/- 24.3 | -20.8 +/- 1.5 | -9.8 +/- 13.7 |
| **Cluster size** | 11 | 4 | 4 | 22 | 4 |
| **RMSD from the overall lowest-energy structure** | 1.1 +/- 0.1 | 2.0 +/- 0.0 | 1.8 +/- 0.1 | 1.5 +/- 0.0 | 1.5 +/- 0.1 |
| **Van der Waals energy** | -55.1 +/- 7.5 | -65.5 +/- 4.2 | -60.1 +/- 9.1 | -43.9 +/- 7.2 | -46.9 +/- 8.5 |
| **Electrostatic energy** | -191.9 +/- 37.6 | -111.9 +/- 17.1 | -72.6 +/- 14.8 | -136.2 +/- 17.2 | -111.9 +/- 26.8 |
| **Desolvation energy** | -1.7 +/- 3.7 | -13.7 +/- 1.2 | -23.2 +/- 5.4 | -5.9 +/- 1.3 | -13.3 +/- 2.8 |
| **Restraints violation energy** | 585.1 +/- 46.8 | 771.0 +/- 58.8 | 749.7 +/- 136.9 | 562.7 +/- 47.4 | 728.5 +/- 132.9 |
| **Buried Surface Area** | 1703.7 +/- 89.5 | 1949.1 +/- 68.5 | 1831.1 +/- 90.0 | 1552.2 +/- 35.9 | 1712.5 +/- 129.7 |
| **Z-Score** | -1.9 | -1.0 | -0.8 | -0.7 | 0.2 |

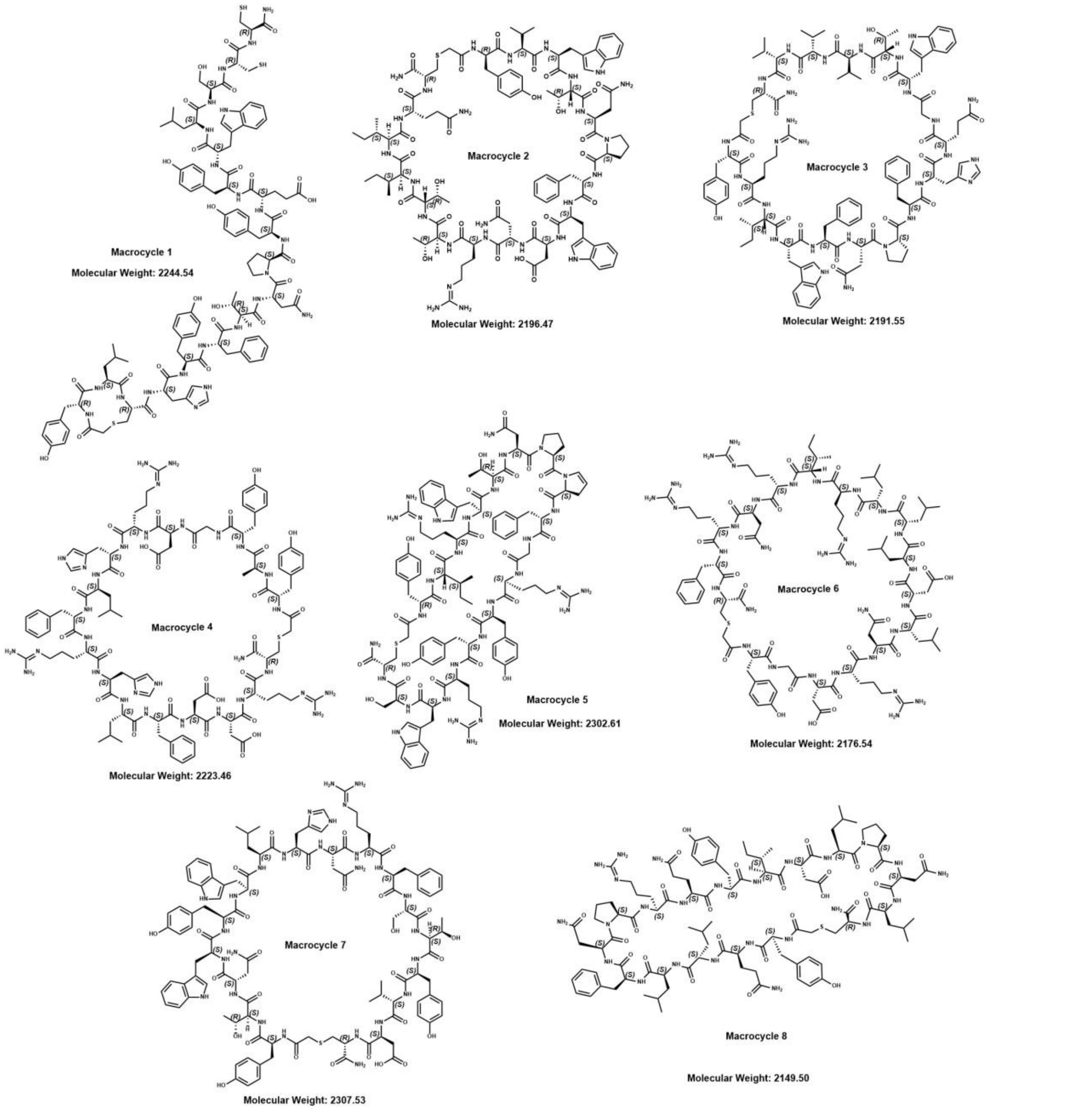


**Supplementary Figure 1.** Eight Macrocycle Structures. Note that 1, 2, and 5 have D-Tyr while 3, 4, 6-8 have L-Tyr. Structures were generated using ChemDraw 23.1.2 (Revvity Signals Software, Inc.).

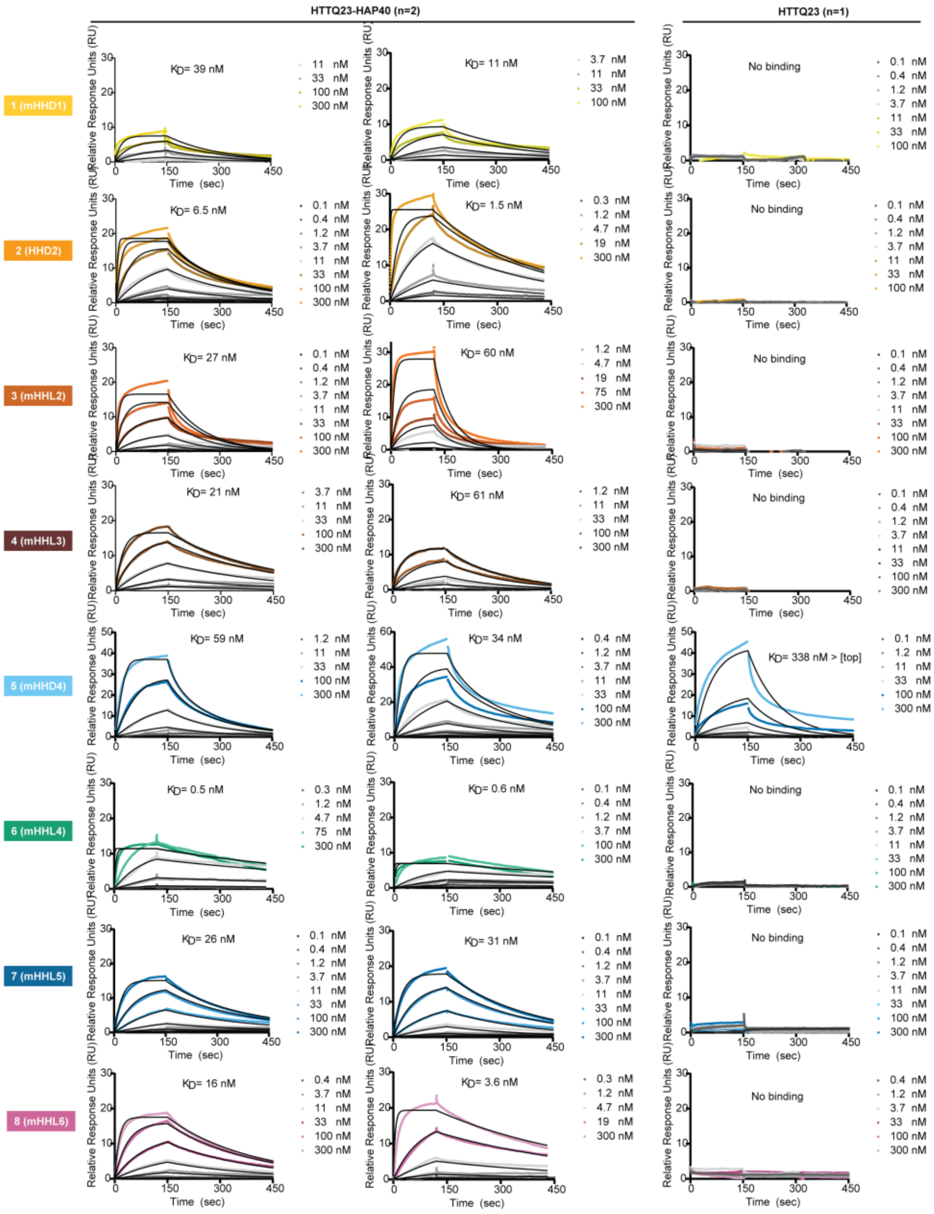


| | HTTQ23-HAP40 (n=2) | | HTTQ23 (n=1) | |
|---|---|---|---|---|
| | Kinetic $K_D$ ± SD (nM) | $Chi^2_{avg}$ ($RU^2$) | Kinetic $K_D$ (nM) | $Chi^2$ ($RU^2$) |
| 1 (mHHD1) | 25 ± 20 | 0.4 | n.b. | n.b. |
| 2 (HHD2) | 4.0 ± 3.5 | 1.0 | n.b. | n.b. |
| 3 (mHHL2) | 44 ± 23 | 1.8 | n.b. | n.b. |
| 4 (mHHL3) | 41 ± 28 | 0.2 | n.b. | n.b. |
| 5 (mHHD4) | 47 ± 18 | 2.7 | 340 | 5.7 |
| 6 (mHHL4) | 0.6 ± 0.1 | 0.8 | n.b. | n.b. |
| 7 (mHHL5) | 29 ± 3.7 | 0.2 | n.b. | n.b. |
| 8 (mHHL6) | 9.8 ± 8.8 | 0.4 | n.b. | n.b. |

**Supplementary Figure 2**. SPR of Eight Macrocycles measured against immobilized HTTQ23 and HTTQ23-HAP40. The summary table lists Kinetic $K_D$ ± Standard Deviation (SD) (nM) of a biological duplicate (n=2) for HTTQ23-HAP40 and biological singlet (n=1) for HTTQ23. n.b., no detectable binding - SPR relative response units did not exceed 8 RU (~20% binding).

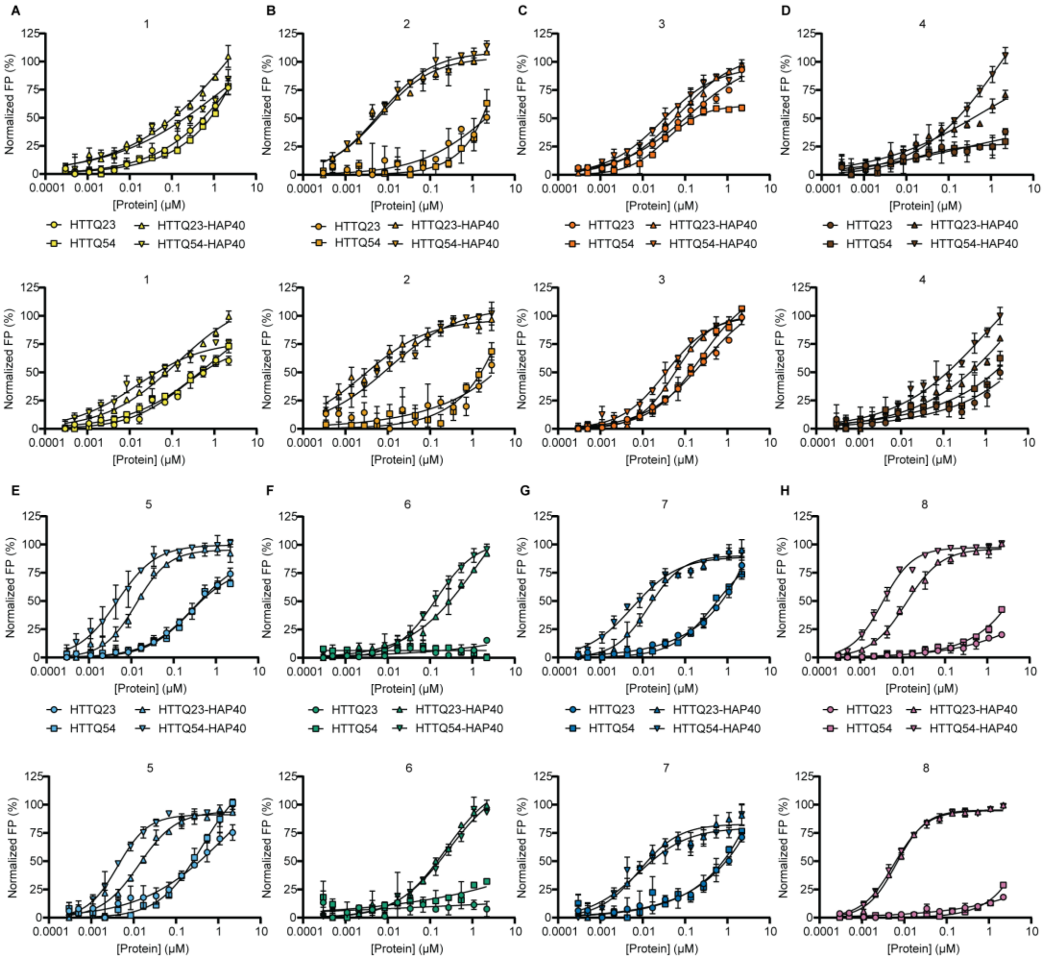


I

| | Full-length: $K_D$ ± SD (nM), n=2 | | | |
|---|---|---|---|---|
| | HTTQ23 | HTTQ54 | HTTQ23-HAP40 | HTTQ54-HAP40 |
| 1 (mHHD1) | n.d. | n.d. | n.d. | n.d. |
| 2 (HHD2) | n.d. | n.d. | 4.3 ± 3.0 | 7.4 ± 0.9 |
| 3 (mHHL2) | n.d. | n.d. | 65 ± 10 | 32 ± 4.5 |
| 4 (mHHL3) | n.d. | n.d. | n.d. | n.d. |
| 5 (mHHD4) | n.d. | n.d. | 13 ± 0.3 | 4.4 ± 0.3 |
| 6 (mHHL4) | n.b. | n.b. | n.d. | n.d. |
| 7 (mHHL5) | n.d. | n.d. | 11 ± 4.1 | 6.9 ± 1.6 |
| 8 (mHHL6) | n.b. | n.b. | 9.3 ± 4.0 | 4.5 ± 1.7 |

**Supplementary Figure 3. HTT and HTT-HAP40 Binding Affinity measured by Fluorescence Polarization. A-H**, Each macrocycle was measured in biological duplicates of technical triplicates against HTTQ23, HTTQ54, HTTQ23-HAP40, and HTTQ54-HAP40. **I**, summary table of $K_D$ ± SD (nM), n=2. n.d., not determined - binding curves not saturated at concentrations tested, $K_D$ exceeds top concentration tested; n.b., no detectable binding - normalized FP did not reach 50% at concentrations tested.

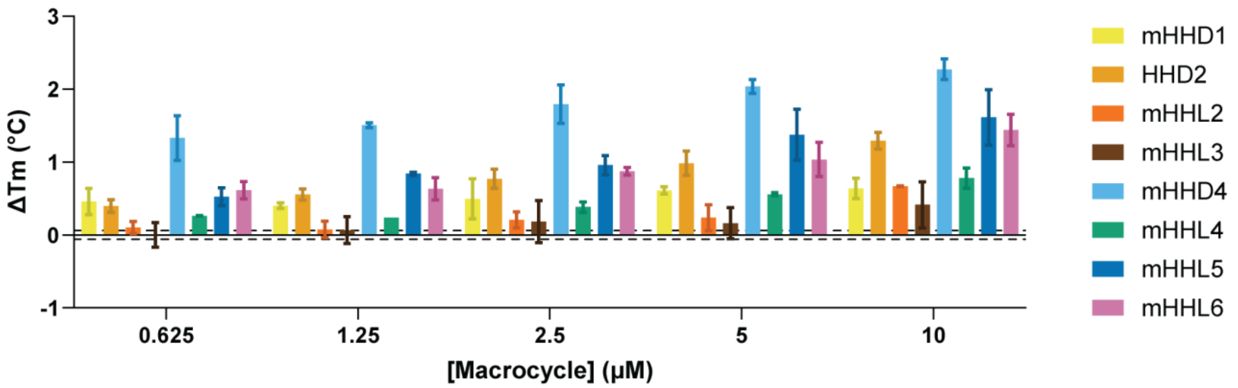


**Supplementary Figure 4.** HTTQ23-HAP40 Thermal Shift Assay (n=2). Thermal stabilization is observed when $\Delta T_m$ exceeds the empirical cut-off of the SD of the HTTQ23-HAP40 DMSO control $T_m$ (dashed line = 0.05 °C).

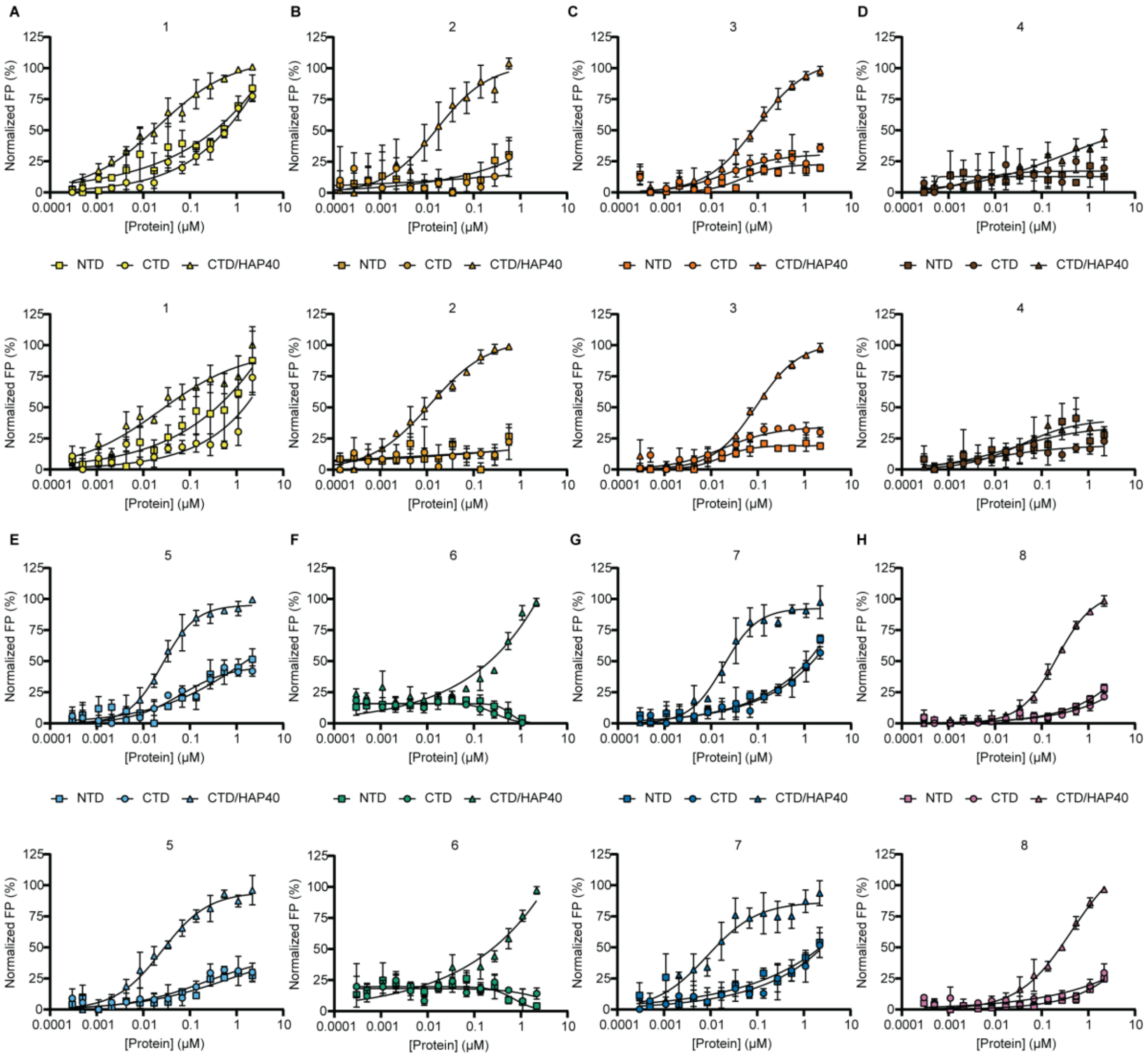


I

| | Subdomains: $K_D$ ± SD (nM), n=2 | | |
|---|---|---|---|
| | NTD | CTD | CTD/HAP40 |
| 1 (mHHD1) | n.d. | n.d. | 23 ± 1.5 |
| 2 (HHD2) | n.b. | n.b. | 14 ± 3.4 |
| 3 (mHHL2) | n.b. | n.b. | 94 ± 1.4 |
| 4 (mHHL3) | n.d. | n.d. | n.d. |
| 5 (mHHD4) | n.d. | n.d. | 25 ± 0.3 |
| 6 (mHHL4) | n.b. | n.b. | n.d. |
| 7 (mHHL5) | n.d. | n.d. | 13 ± 5.9 |
| 8 (mHHL6) | n.b. | n.b. | 330 ± 160 |

**Supplementary Figure 5.** Subdomain Binding Affinity measured by Fluorescence Polarization. **A-H**, Each macrocycle was measured in biological duplicates of technical triplicates against NTD, CTD, CTD/HAP40. **I**, summary table of $K_D$ ± SD (nM), n=2. Binding was not determined (n.d.) if curves did not saturate at concentrations tested. Binding was not detected (n.b.) if normalized FP did not reach 50% at concentrations tested.

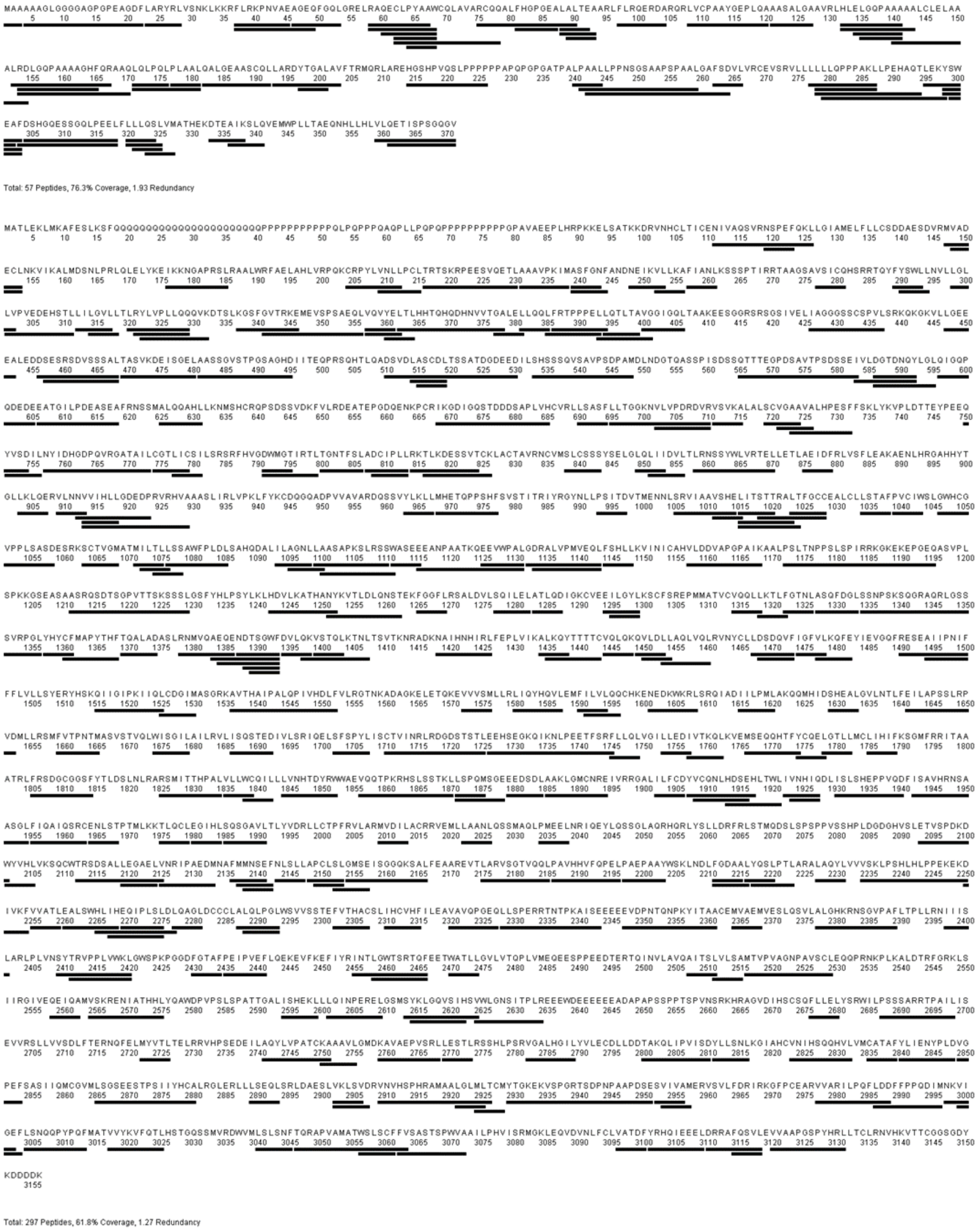


**Supplementary Figure 6.** HDX-MS HTTQ23-HAP40 Peptide Sequence Coverage for Macrocycles 1,3-6.

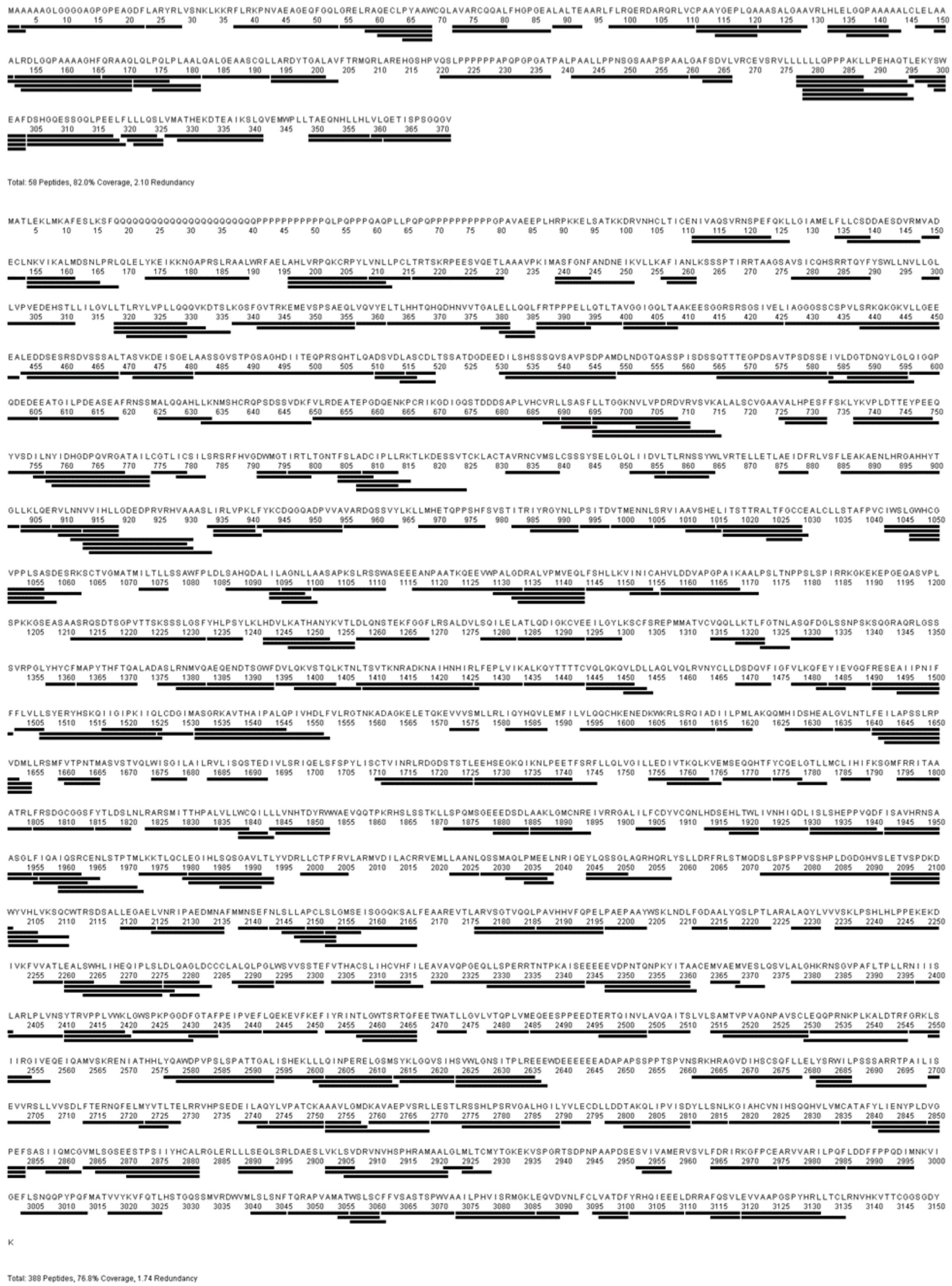


**Supplementary Figure 7.** HDX-MS HTTQ23-HAP40 Peptide Sequence Coverage for Macrocycles 2.

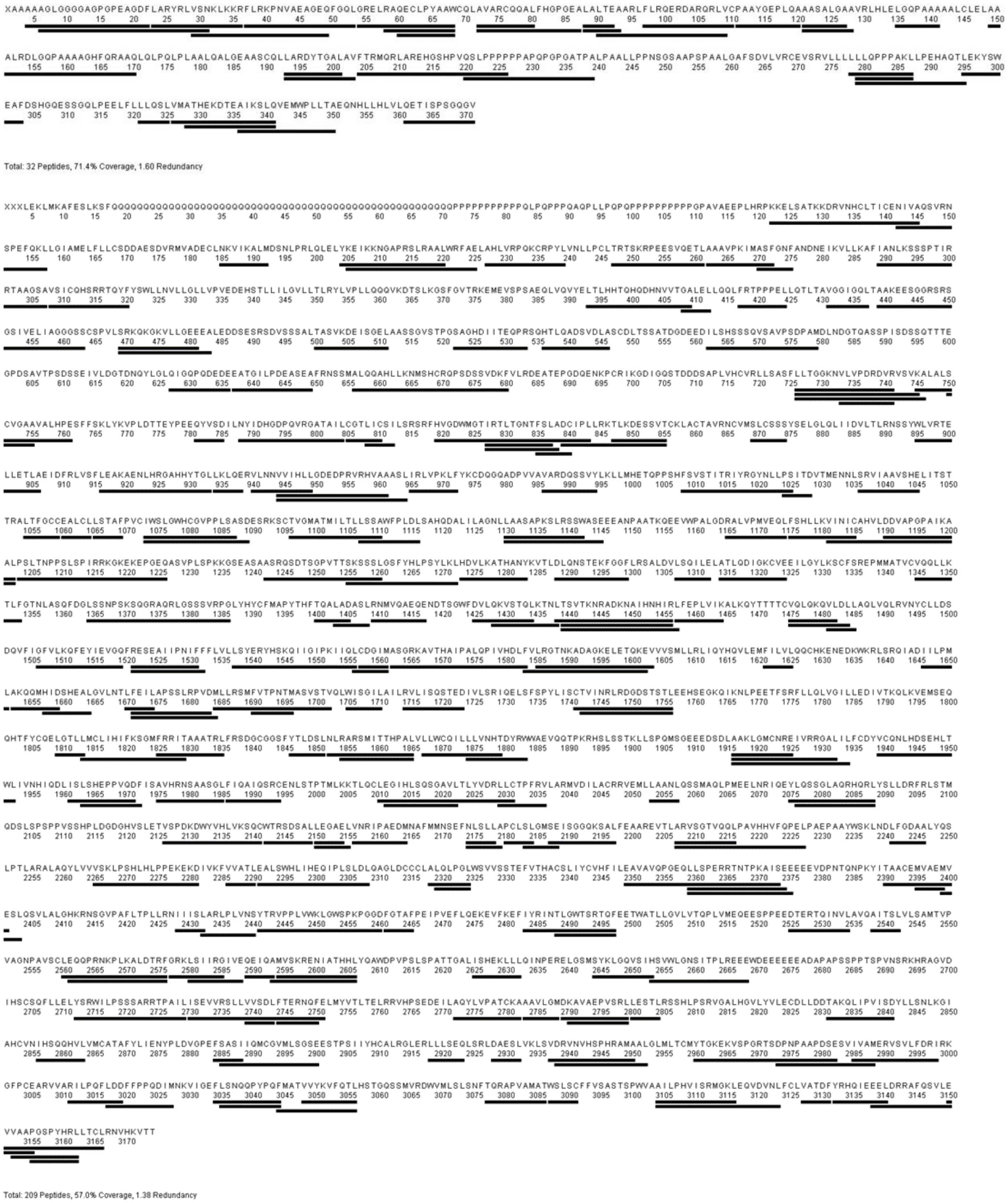


**Supplementary Figure 8.** HDX-MS HTTQ54-HAP40 Peptide Sequence Coverage for Macrocycles 7 and 8.

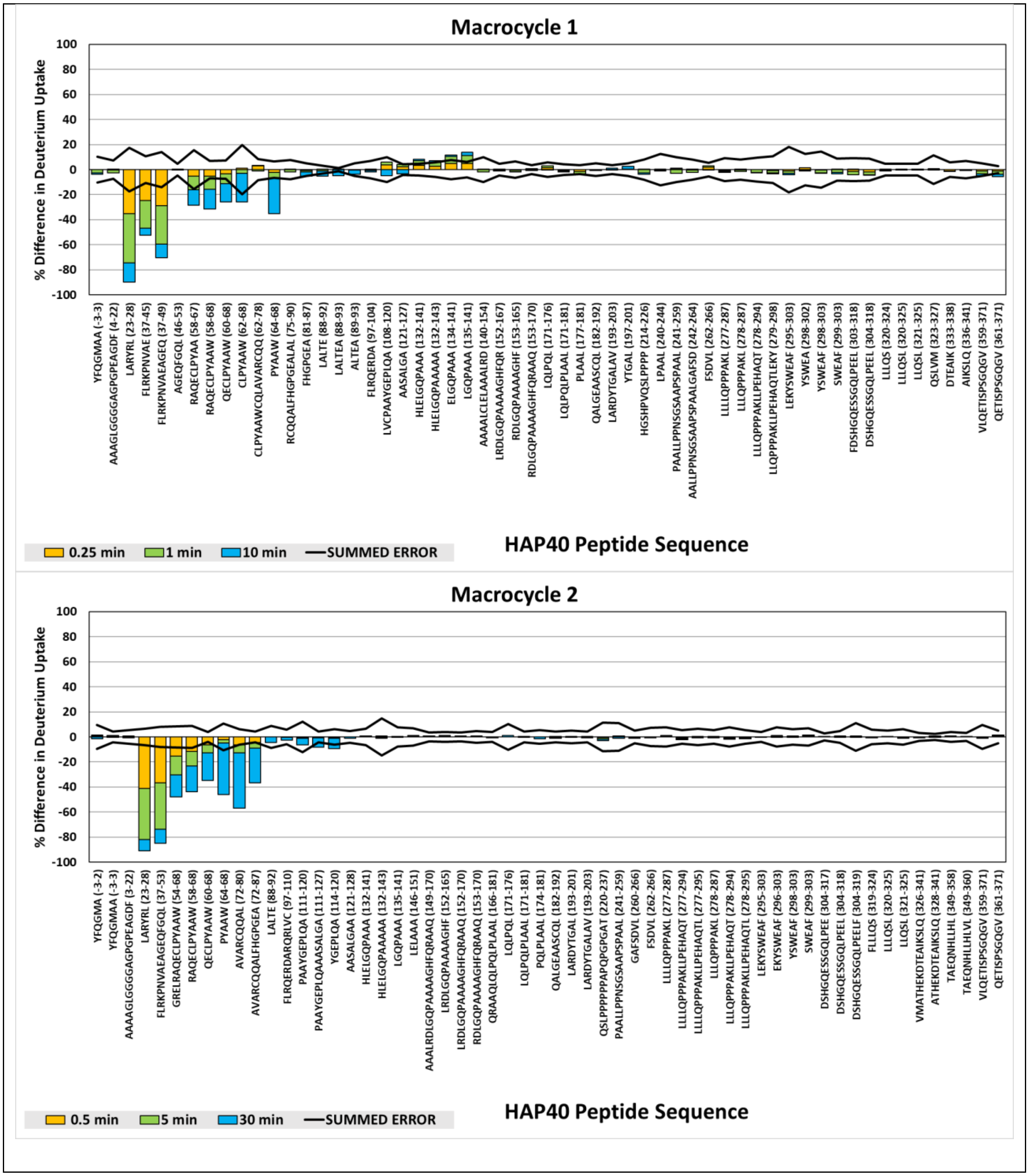
Macrocycle 1
% Difference in Deuterium Uptake
100
80
60
40
20
0
-20
-40
-60
-80
-100
YFQGMAA (-3-3)
AAAGLGGGGAGPGPEAGDF (4-22)
LARYRL (23-28)
FLRKPNVAE (37-45)
FLRKPNVAEAGEQ (37-49)
AGEQFGQL (46-53)
RAQECLPYAA (58-67)
RAQECLPYAAW (58-68)
QECLPYAAW (60-68)
CLPYAAW (62-68)
CLPYAAWCQLAVARCQQ (62-78)
PYAAW (64-68)
RCQQALFHGPGEALAL (75-90)
FHGPGEA (81-87)
LALTE (88-92)
LALTEA (88-93)
ALTEA (89-93)
FLRQERDA (97-104)
LVCPAAYGEPLQA (108-120)
AASALGA (121-127)
HLELGQPAAA (132-141)
HLELGQPAAAA (132-143)
ELGQPAAA (134-141)
LGQPAAA (135-141)
AAAALCLELAAALRD (140-154)
LRDLGQPAAAAGHFQR (152-167)
RDLGQPAAAAGHF (153-165)
RDLGQPAAAAGHFQRAAQ (153-170)
LQLPQL (171-176)
LQLPQLPLAAL (171-181)
PLAAL (177-181)
QALGEAASCQL (182-192)
LARDYTGALAV (193-203)
YTGAL (197-201)
HGSHPVQSLPPPP (214-226)
LPAAL (240-244)
PAALLPPNSGSAAPSPAAL (241-259)
AALLPPNSGSAAPSPAALGAFSD (242-264)
FSDVL (262-266)
LLLLQPPPAKL (277-287)
LLLQPPAKL (278-287)
LLLQPPPAKLLPEHAQT (278-294)
LLQPPPAKLLPEHAQTLEKY (279-298)
LEKYSWEAF (295-303)
YSWEA (298-302)
YSWEAF (298-303)
SWEAF (299-303)
FDSHGQESSGQLPEEL (303-318)
DSHGQESSGQLPEEL (304-318)
LLLQS (320-324)
LLLQSL (320-325)
LLQSL (321-325)
QSLVM (323-327)
DTEAIK (333-338)
AIKSLQ (336-341)
VLQETISPSGQGV (359-371)
QETISPSGQGV (361-371)
0.25 min
1 min
10 min
SUMMED ERROR
HAP40 Peptide Sequence
Macrocycle 2
% Difference in Deuterium Uptake
100
80
60
40
20
0
-20
-40
-60
-80
-100
YFQGMA (-3-2)
YFQGMAA (-3-3)
AAAGLGGGGAGPGPEAGDF (3-22)
LARYRL (23-28)
FLRKPNVAEAGEQFGQL (37-53)
GRELRAQECLPYAAW (54-68)
RAQECLPYAAW (58-68)
QECLPYAAW (60-68)
PYAAW (64-68)
AVARCQQAL (72-80)
AVARCQQALFHGPGEA (72-87)
LALTE (88-92)
FLRQERDARQRLVC (97-110)
PAAYGEPLQA (111-120)
PAAYGEPLQAAASALGA (111-127)
YGEPLQA (114-120)
AASALGAA (121-128)
HLELGQPAAA (132-141)
HLELGQPAAAA (132-143)
LGQPAAA (135-141)
LELAAA (146-151)
AAALRDLGQPAAAAGHFQRAAQ (149-170)
LRDLGQPAAAAGHF (152-165)
LRDLGQPAAAAGHFQRAAQ (152-170)
RDLGQPAAAAGHFQRAAQ (153-170)
QRAAQLQLPQLPLAAL (166-181)
LQLPQL (171-176)
LQLPQLPLAAL (171-181)
PQLPLAAL (174-181)
QALGEAASCQL (182-192)
LARDYTGAL (193-201)
LARDYTGALAV (193-203)
QSLPPPPAPQPGPGAT (220-237)
PAALLPPNSGSAAPSPAAL (241-259)
GAFSDVL (260-266)
FSDVL (262-266)
LLLLQPPPAKL (277-287)
LLLLQPPPAKLLPEHAQT (277-294)
LLLLQPPPAKLLPEHAQTL (277-295)
LLLQPPPAKL (278-287)
LLLQPPPAKLLPEHAQT (278-294)
LLLQPPPAKLLPEHAQTL (278-295)
LEKYSWEAF (295-303)
EKYSWEAF (296-303)
YSWEAF (298-303)
SWEAF (299-303)
DSHGQESSGQLPEE (304-317)
DSHGQESSGQLPEEL (304-318)
DSHGQESSGQLPEELF (304-319)
FLLLQS (319-324)
LLLQSL (320-325)
LLQSL (321-325)
VMATHEKDTEAIKSLQ (326-341)
ATHEKDTEAIKSLQ (328-341)
TAEQNHLLHL (349-358)
TAEQNHLLHLVL (349-360)
VLQETISPSGQGV (359-371)
QETISPSGQGV (361-371)
0.5 min
5 min
30 min
SUMMED ERROR
HAP40 Peptide Sequence

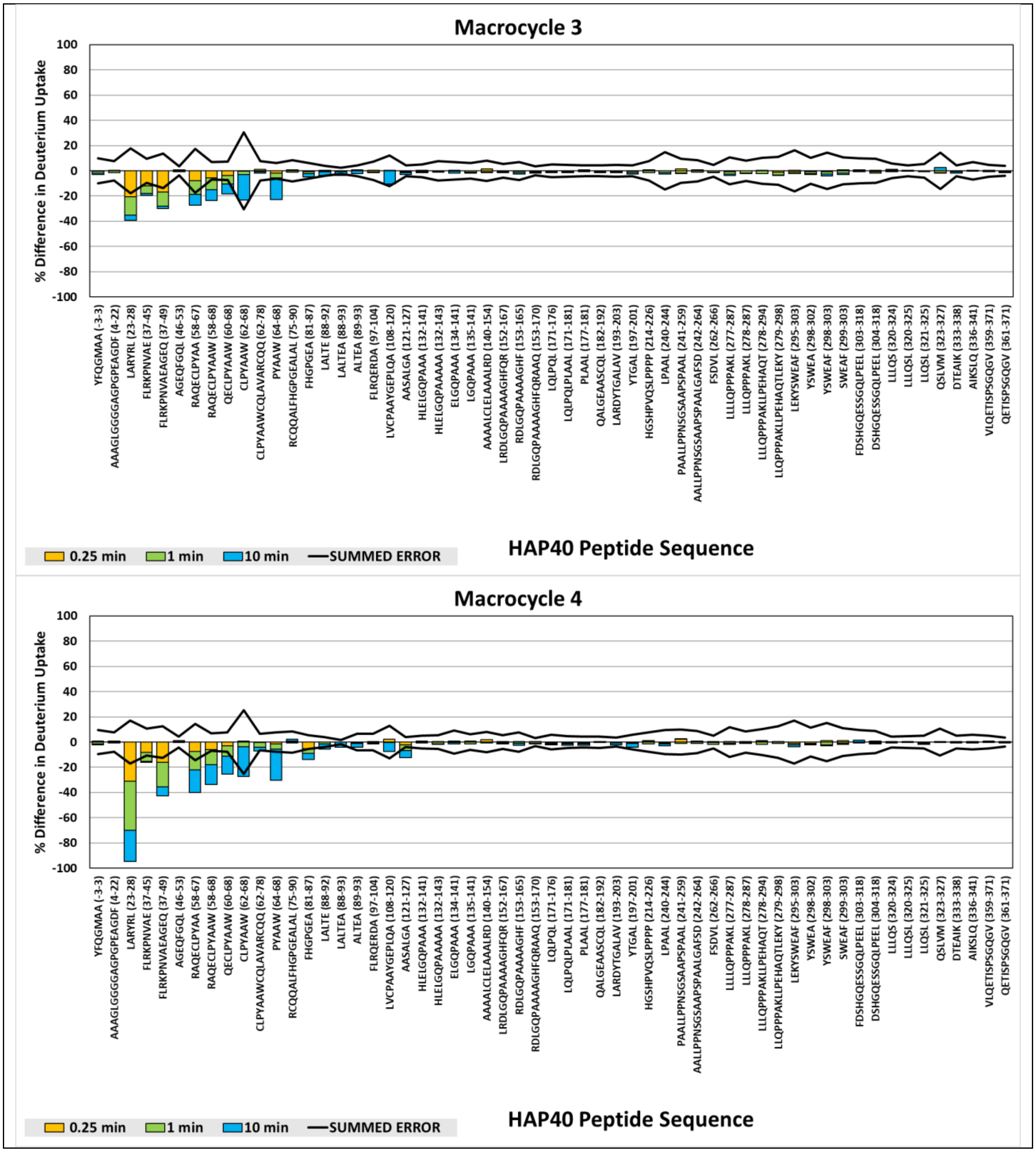
Macrocycle 3
% Difference in Deuterium Uptake
100
80
60
40
20
0
-20
-40
-60
-80
-100
YFQGMAA (-3-3)
AAAGLGGGGAGPGPEAGDF (4-22)
LARYRL (23-28)
FLRKPNVAE (37-45)
FLRKPNVAEAGEQ (37-49)
AGEQFGQL (46-53)
RAQECLPYAA (58-67)
RAQECLPYAAW (58-68)
QECLPYAAW (60-68)
CLPYAAW (62-68)
CLPYAAWCQLAVARCQQ (62-78)
PYAAW (64-68)
RCQQALFHGPGEALAL (75-90)
FHGPGEA (81-87)
LALTE (88-92)
LALTEA (88-93)
ALTEA (89-93)
FLRQERDA (97-104)
LVCPAAYGEPLQA (108-120)
AASALGA (121-127)
HLEIGQPAAA (132-141)
HLEIGQPAAAAA (132-143)
ELGQPAAA (134-141)
LGQPAAA (135-141)
AAAALCLELAAALRD (140-154)
LRDLGQPAAAAGHFQR (152-167)
RDLGQPAAAAGHF (153-165)
RDLGQPAAAAGHFQRAAQ (153-170)
LQLPQL (171-176)
LQLPQLPLAAL (171-181)
PLAAL (177-181)
QALGEAASCQL (182-192)
LARDYTGALAV (193-203)
YTGAL (197-201)
HGSHPVQSLPPPP (214-226)
LPAAL (240-244)
PAALLPPNSGSAAPSPAAL (241-259)
AALLPPNSGSAAPSPAALGAFSD (242-264)
FSDVL (262-266)
LLLLQPPPAKL (277-287)
LLLQPPPAKL (278-287)
LLLQPPPAKLLPEHAQT (278-294)
LLQPPPAKLLPEHAQTLEKY (279-298)
LEKYSWEAF (295-303)
YSWEA (298-302)
YSWEAF (298-303)
SWEAF (299-303)
FDSHGQESSGQLPEEL (303-318)
DSHGQESSGQLPEEL (304-318)
LLLQS (320-324)
LLLQSL (320-325)
LLQSL (321-325)
QSLVM (323-327)
DTEAIK (333-338)
AIKSLQ (336-341)
VLQETISPSGQGV (359-371)
QETISPSGQGV (361-371)
0.25 min
1 min
10 min
SUMMED ERROR
HAP40 Peptide Sequence
Macrocycle 4
% Difference in Deuterium Uptake
100
80
60
40
20
0
-20
-40
-60
-80
-100
YFQGMAA (-3-3)
AAAGLGGGGAGPGPEAGDF (4-22)
LARYRL (23-28)
FLRKPNVAE (37-45)
FLRKPNVAEAGEQ (37-49)
AGEQFGQL (46-53)
RAQECLPYAA (58-67)
RAQECLPYAAW (58-68)
QECLPYAAW (60-68)
CLPYAAW (62-68)
CLPYAAWCQLAVARCQQ (62-78)
PYAAW (64-68)
RCQQALFHGPGEALAL (75-90)
FHGPGEA (81-87)
LALTE (88-92)
LALTEA (88-93)
ALTEA (89-93)
FLRQERDA (97-104)
LVCPAAYGEPLQA (108-120)
AASALGA (121-127)
HLEIGQPAAA (132-141)
HLEIGQPAAAAA (132-143)
ELGQPAAA (134-141)
LGQPAAA (135-141)
AAAALCLELAAALRD (140-154)
LRDLGQPAAAAGHFQR (152-167)
RDLGQPAAAAGHF (153-165)
RDLGQPAAAAGHFQRAAQ (153-170)
LQLPQL (171-176)
LQLPQLPLAAL (171-181)
PLAAL (177-181)
QALGEAASCQL (182-192)
LARDYTGALAV (193-203)
YTGAL (197-201)
HGSHPVQSLPPPP (214-226)
LPAAL (240-244)
PAALLPPNSGSAAPSPAAL (241-259)
AALLPPNSGSAAPSPAALGAFSD (242-264)
FSDVL (262-266)
LLLLQPPPAKL (277-287)
LLLQPPPAKL (278-287)
LLLQPPPAKLLPEHAQT (278-294)
LLQPPPAKLLPEHAQTLEKY (279-298)
LEKYSWEAF (295-303)
YSWEA (298-302)
YSWEAF (298-303)
SWEAF (299-303)
FDSHGQESSGQLPEEL (303-318)
DSHGQESSGQLPEEL (304-318)
LLLQS (320-324)
LLLQSL (320-325)
LLQSL (321-325)
QSLVM (323-327)
DTEAIK (333-338)
AIKSLQ (336-341)
VLQETISPSGQGV (359-371)
QETISPSGQGV (361-371)
0.25 min
1 min
10 min
SUMMED ERROR
HAP40 Peptide Sequence

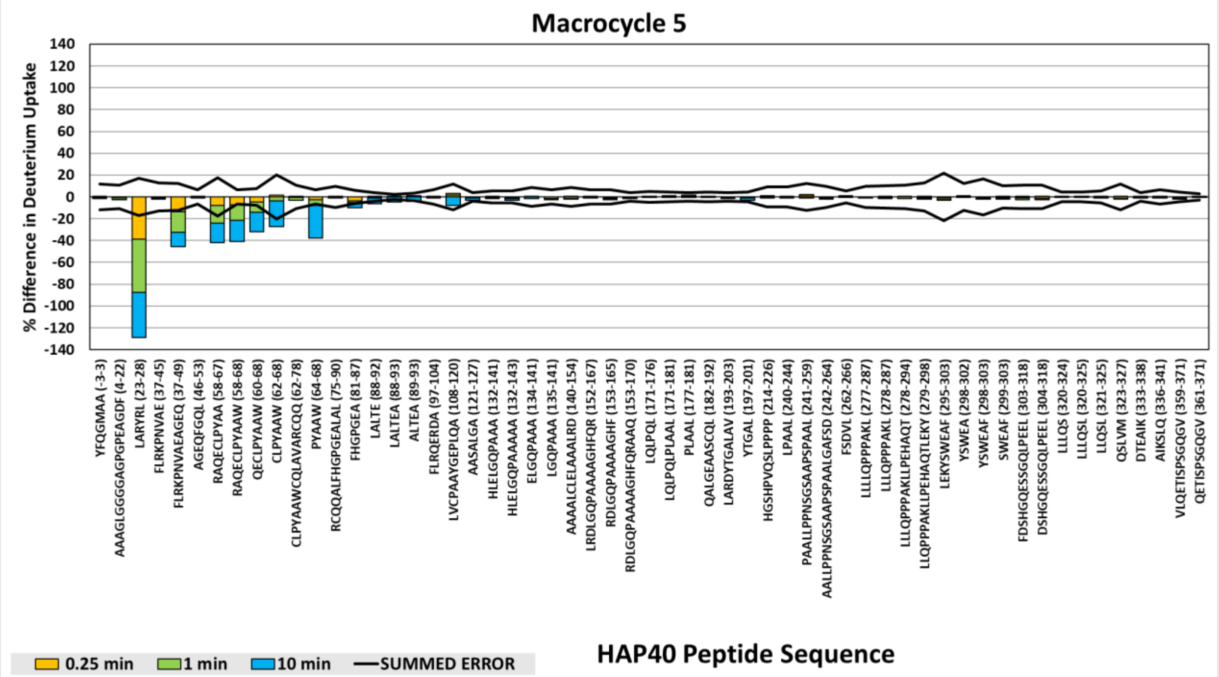
Macrocycle 5
% Difference in Deuterium Uptake
140
120
100
80
60
40
20
0
-20
-40
-60
-80
-100
-120
-140
YFQGMAA (-3-3)
AAAGLGGGGAGPGPEAGDF (4-22)
LARYRL (23-28)
FLRKPNVAE (37-45)
FLRKPNVAEAGEQ (37-49)
AGEQFGQL (46-53)
RAQECLPYAA (58-67)
RAQECLPYAAW (58-68)
QECLPYAAW (60-68)
CLPYAAW (62-68)
CLPYAAWCQLAVARCQQ (62-78)
PYAAW (64-68)
RCQQALFHGPGEALAL (75-90)
FHGPGEA (81-87)
LALTE (88-92)
LALTEA (88-93)
ALTEA (89-93)
FLRQERDA (97-104)
LVCPAAYGEPLQA (108-120)
AASALGA (121-127)
HLELGQPAAA (132-141)
HLELGQPAAAAA (132-143)
ELGQPAAA (134-141)
LGQPAAA (135-141)
AAAALCLELAAALRD (140-154)
LRDLGQPAAAAGHFQR (152-167)
RDLGQPAAAAGHF (153-165)
RDLGQPAAAAGHFQRAAQ (153-170)
LQLPQL (171-176)
LQLPQLPLAAL (171-181)
PLAAL (177-181)
QALGEAASCQL (182-192)
LARDYTGALAV (193-203)
YTGAL (197-201)
HGSHPVQSLPPPP (214-226)
LPAAL (240-244)
PAALLPPNSGSAAPSPAAL (241-259)
AALLPPNSGSAAPSPAALGAFSD (242-264)
FSDVL (262-266)
LLLLQPPPAKL (277-287)
LLLQPPPAKL (278-287)
LLLQPPPAKLLPEHAQT (278-294)
LLQPPPAKLLPEHAQTLEKY (279-298)
LEKYSWEAF (295-303)
YSWEA (298-302)
YSWEAF (298-303)
SWEAF (299-303)
FDSHGQESSGQLPEEL (303-318)
DSHGQESSGQLPEEL (304-318)
LLLQS (320-324)
LLLQSL (320-325)
LLQSL (321-325)
QSLVM (323-327)
DTEAIK (333-338)
AIKSLQ (336-341)
VLQETISPSGQGV (359-371)
QETISPSGQGV (361-371)
0.25 min
1 min
10 min
SUMMED ERROR
HAP40 Peptide Sequence

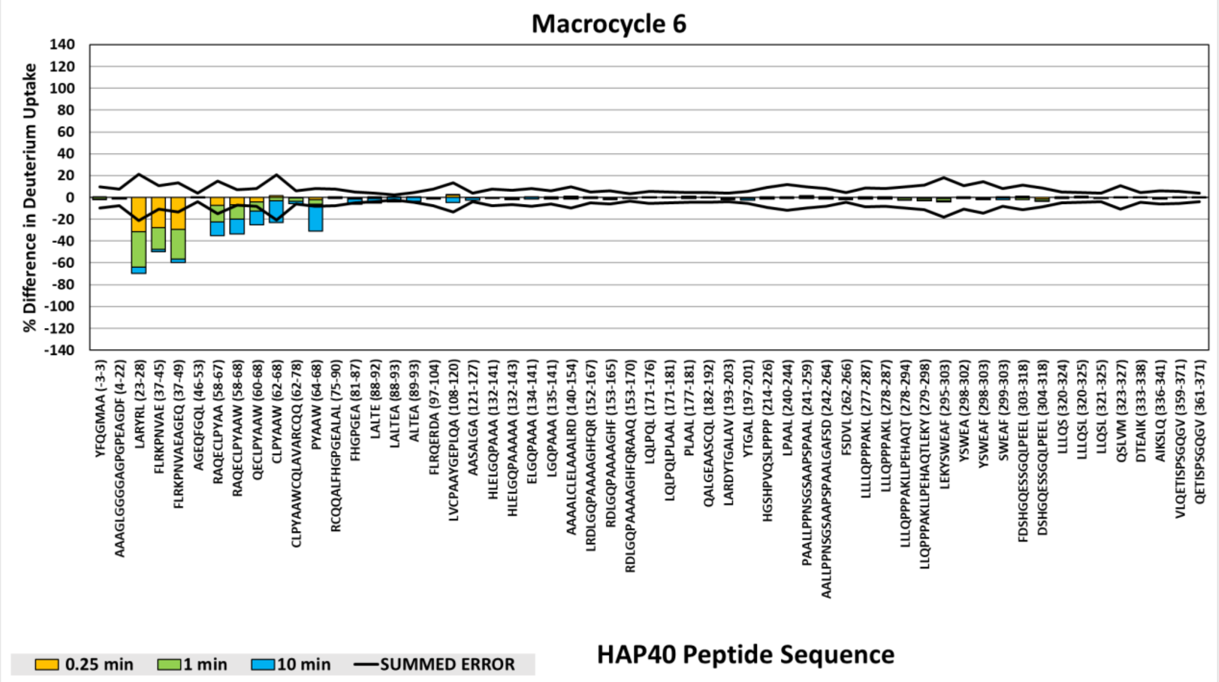
Macrocycle 6
% Difference in Deuterium Uptake
140
120
100
80
60
40
20
0
-20
-40
-60
-80
-100
-120
-140
YFQGMAA (-3-3)
AAAGLGGGGAGPGPEAGDF (4-22)
LARYRL (23-28)
FLRKPNVAE (37-45)
FLRKPNVAEAGEQ (37-49)
AGEQFGQL (46-53)
RAQECLPYAA (58-67)
RAQECLPYAAW (58-68)
QECLPYAAW (60-68)
CLPYAAW (62-68)
CLPYAAWCQLAVARCQQ (62-78)
PYAAW (64-68)
RCQQALFHGPGEALAL (75-90)
FHGPGEA (81-87)
LALTE (88-92)
LALTEA (88-93)
ALTEA (89-93)
FLRQERDA (97-104)
LVCPAAYGEPLQA (108-120)
AASALGA (121-127)
HLELGQPAAA (132-141)
HLELGQPAAAAA (132-143)
ELGQPAAA (134-141)
LGQPAAA (135-141)
AAAALCLELAAALRD (140-154)
LRDLGQPAAAAGHFQR (152-167)
RDLGQPAAAAGHF (153-165)
RDLGQPAAAAGHFQRAAQ (153-170)
LQLPQL (171-176)
LQLPQLPLAAL (171-181)
PLAAL (177-181)
QALGEAASCQL (182-192)
LARDYTGALAV (193-203)
YTGAL (197-201)
HGSHPVQSLPPPP (214-226)
LPAAL (240-244)
PAALLPPNSGSAAPSPAAL (241-259)
AALLPPNSGSAAPSPAALGAFSD (242-264)
FSDVL (262-266)
LLLLQPPPAKL (277-287)
LLLQPPPAKL (278-287)
LLLQPPPAKLLPEHAQT (278-294)
LLQPPPAKLLPEHAQTLEKY (279-298)
LEKYSWEAF (295-303)
YSWEA (298-302)
YSWEAF (298-303)
SWEAF (299-303)
FDSHGQESSGQLPEEL (303-318)
DSHGQESSGQLPEEL (304-318)
LLLQS (320-324)
LLLQSL (320-325)
LLQSL (321-325)
QSLVM (323-327)
DTEAIK (333-338)
AIKSLQ (336-341)
VLQETISPSGQGV (359-371)
QETISPSGQGV (361-371)
0.25 min
1 min
10 min
SUMMED ERROR
HAP40 Peptide Sequence

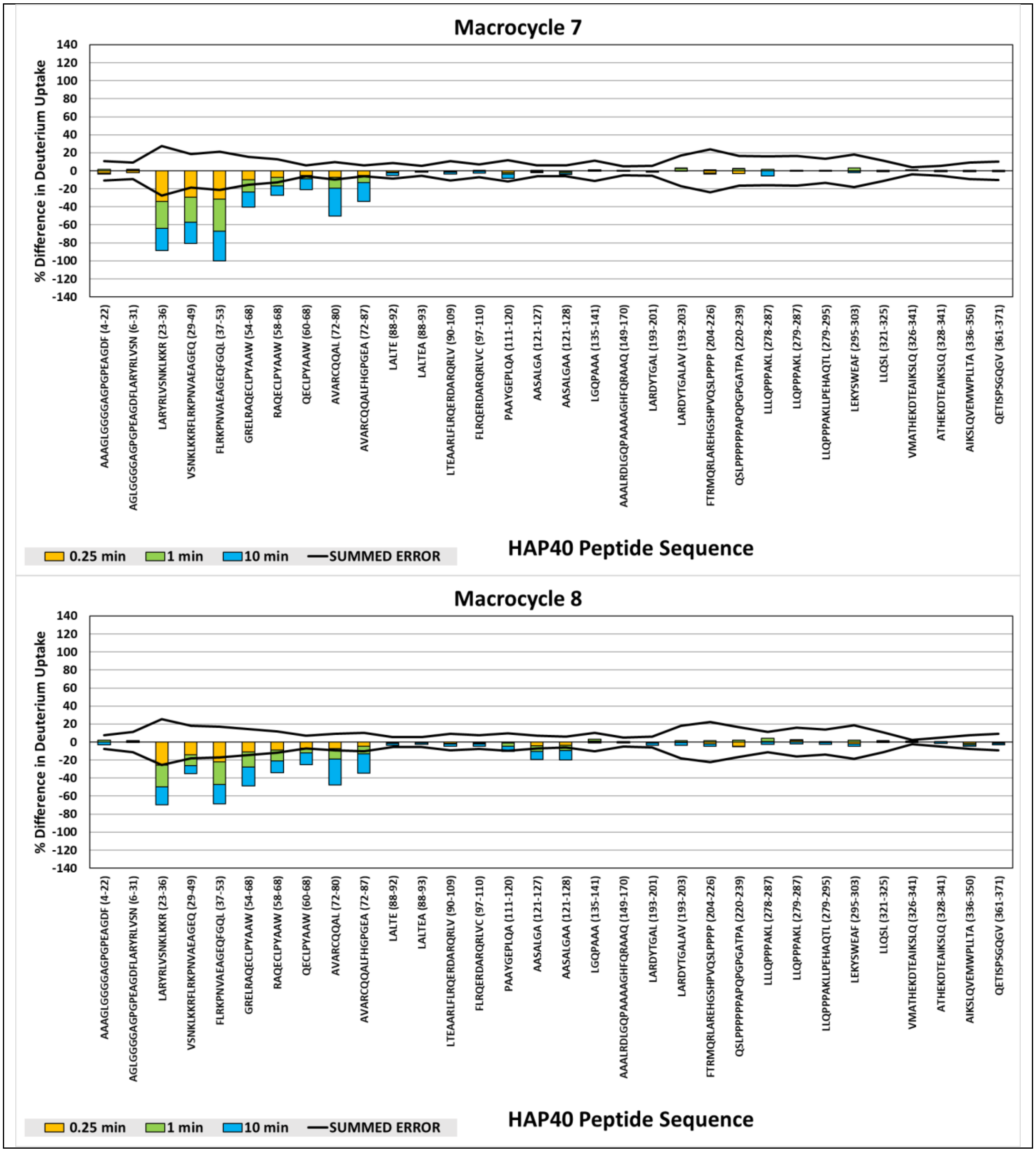


**Supplementary Figure 9.** HAP40 Timepoint-Level ΔHDX-MS.

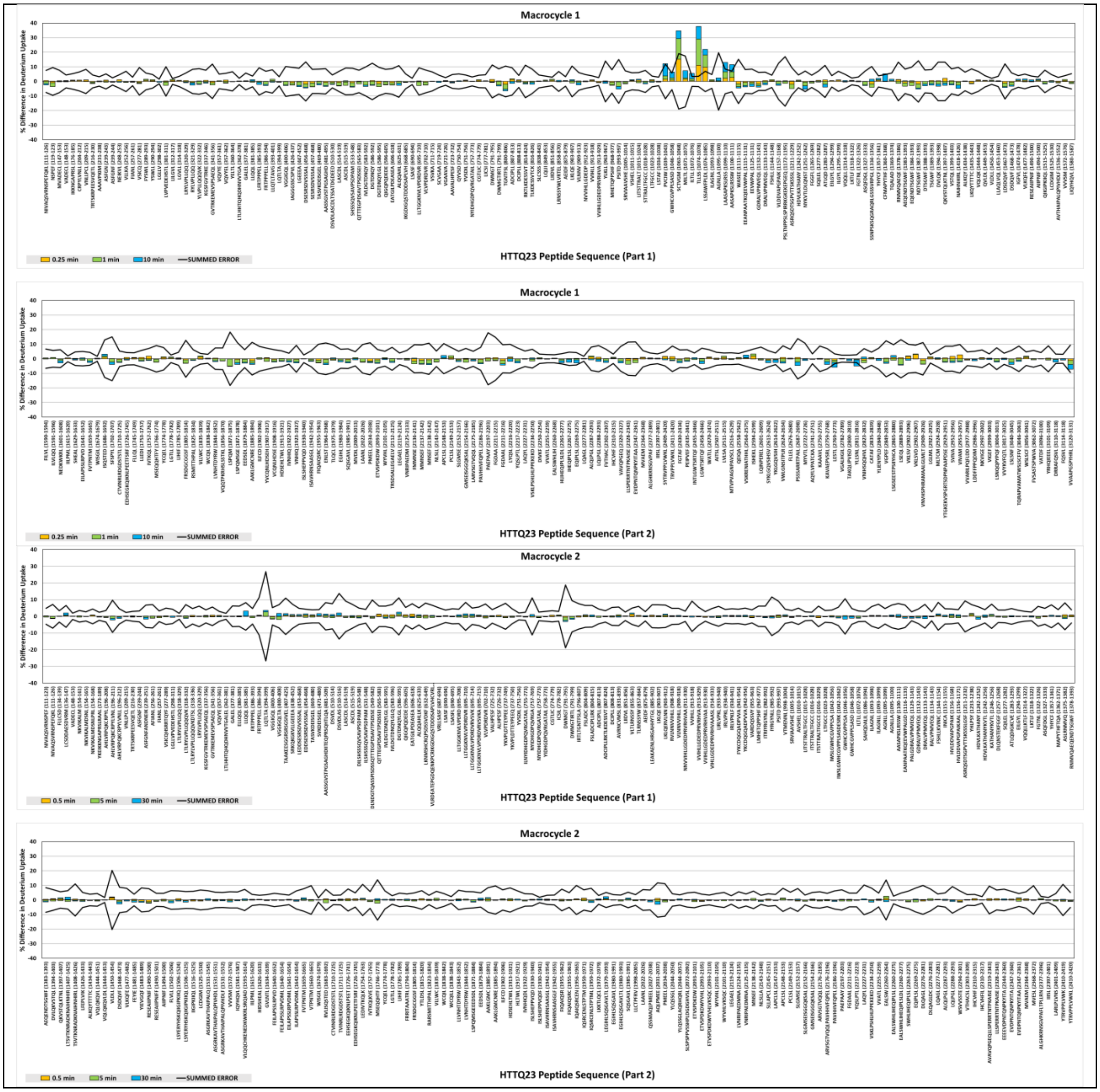
Macrocycle 1
% Difference in Deuterium Uptake
HTTQ23 Peptide Sequence (Part 1)
0.25 min
1 min
10 min
SUMMED ERROR
Macrocycle 1
% Difference in Deuterium Uptake
HTTQ23 Peptide Sequence (Part 2)
0.25 min
1 min
10 min
SUMMED ERROR
Macrocycle 2
% Difference in Deuterium Uptake
HTTQ23 Peptide Sequence (Part 1)
0.5 min
5 min
30 min
SUMMED ERROR
Macrocycle 2
% Difference in Deuterium Uptake
HTTQ23 Peptide Sequence (Part 2)
0.5 min
5 min
30 min
SUMMED ERROR

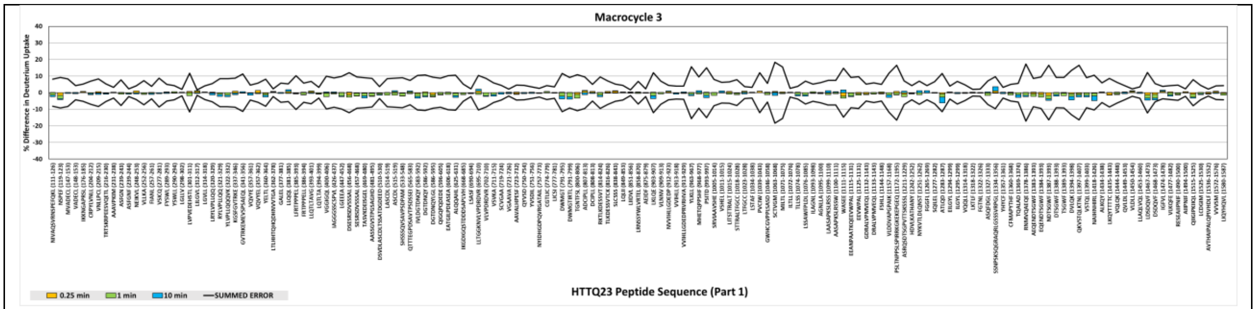
Macrocycle 3
% Difference in Deuterium Uptake
40
30
20
10
0
-10
-20
-30
-40
0.25 min
1 min
10 min
SUMMED ERROR
HTTQ23 Peptide Sequence (Part 1)

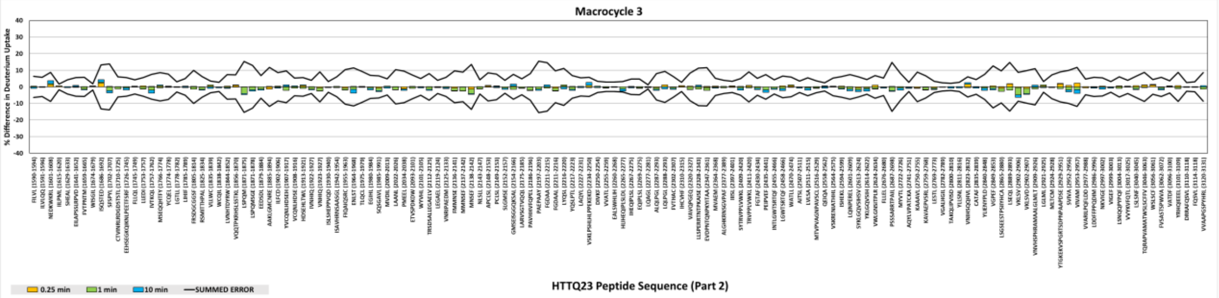
Macrocycle 3
% Difference in Deuterium Uptake
40
30
20
10
0
-10
-20
-30
-40
0.25 min
1 min
10 min
SUMMED ERROR
HTTQ23 Peptide Sequence (Part 2)

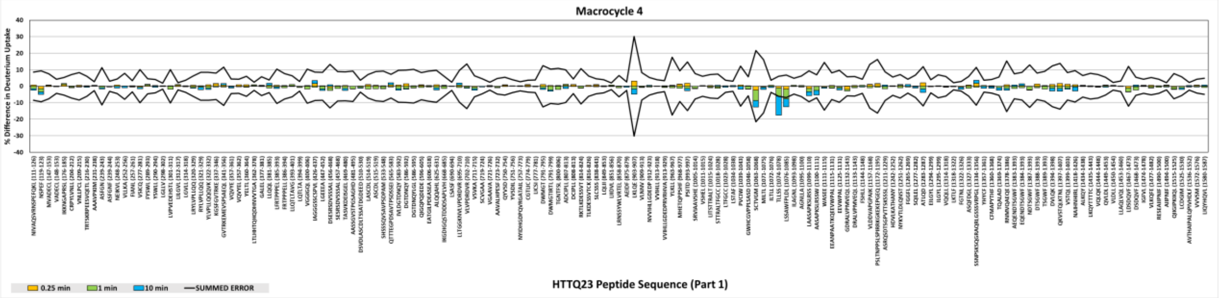
Macrocycle 4
% Difference in Deuterium Uptake
40
30
20
10
0
-10
-20
-30
-40
0.25 min
1 min
10 min
SUMMED ERROR
HTTQ23 Peptide Sequence (Part 1)

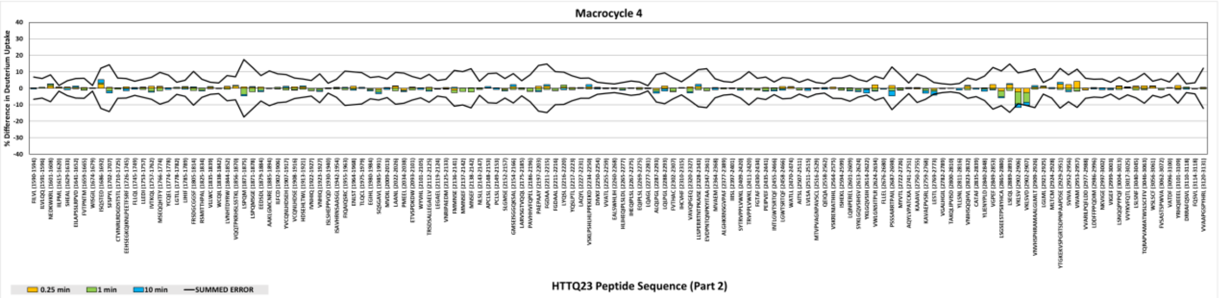
Macrocycle 4
% Difference in Deuterium Uptake
40
30
20
10
0
-10
-20
-30
-40
0.25 min
1 min
10 min
SUMMED ERROR
HTTQ23 Peptide Sequence (Part 2)

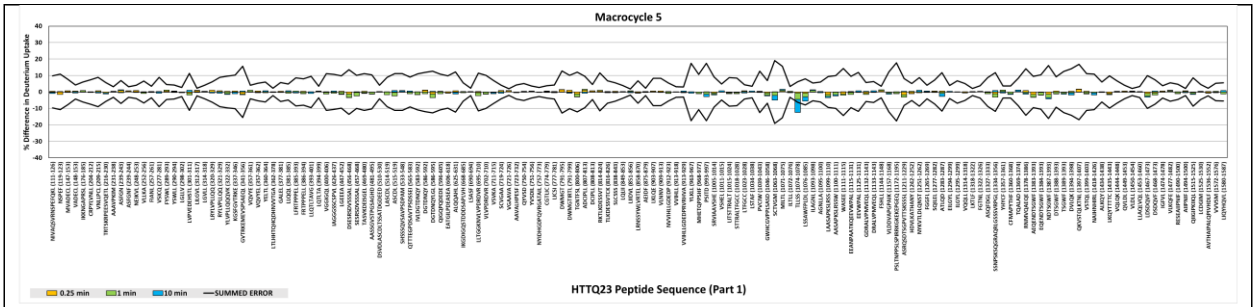

Macrocycle 5
% Difference in Deuterium Uptake
HTTQ23 Peptide Sequence (Part 1)
0.25 min
1 min
10 min
SUMMED ERROR


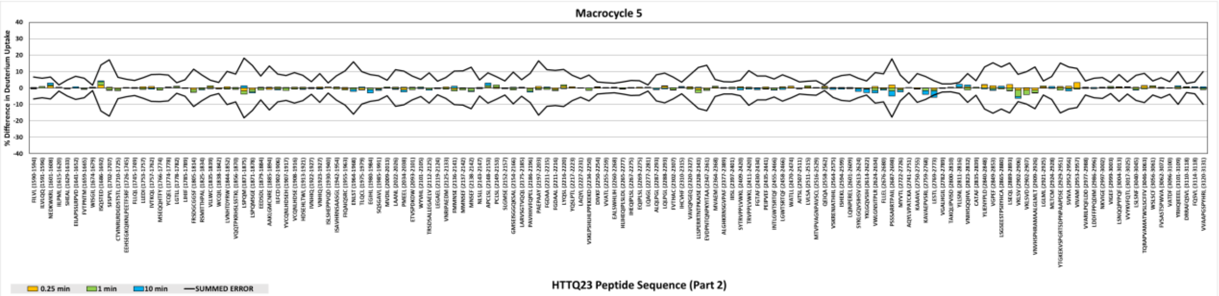

Macrocycle 5
% Difference in Deuterium Uptake
HTTQ23 Peptide Sequence (Part 2)
0.25 min
1 min
10 min
SUMMED ERROR


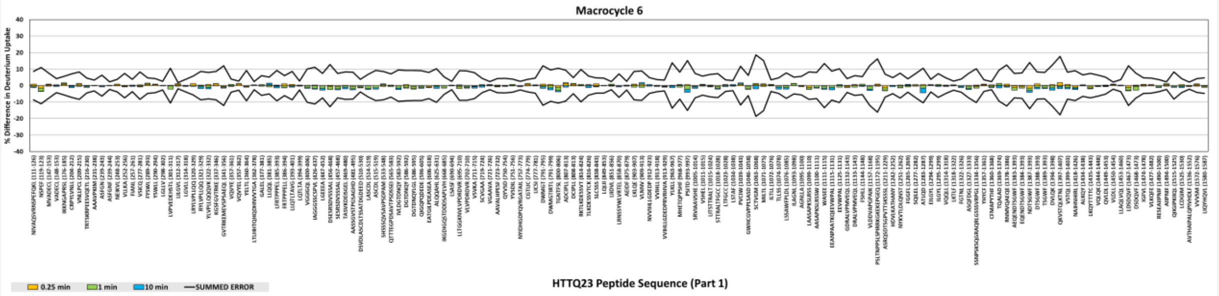

Macrocycle 6
% Difference in Deuterium Uptake
HTTQ23 Peptide Sequence (Part 1)
0.25 min
1 min
10 min
SUMMED ERROR


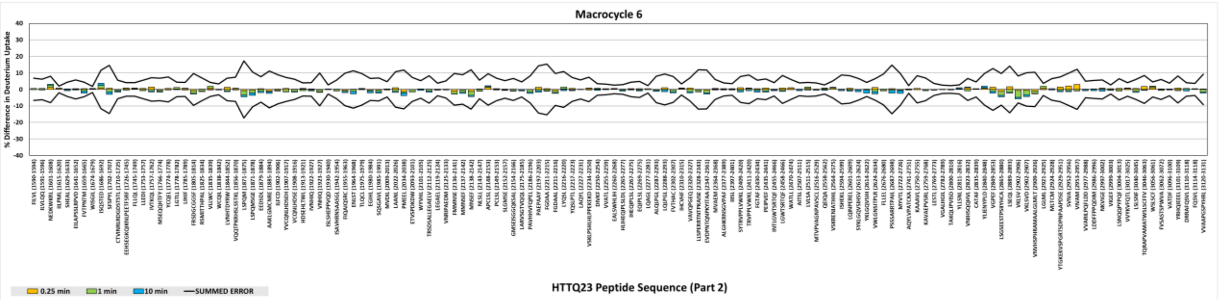

Macrocycle 6
% Difference in Deuterium Uptake
HTTQ23 Peptide Sequence (Part 2)
0.25 min
1 min
10 min
SUMMED ERROR

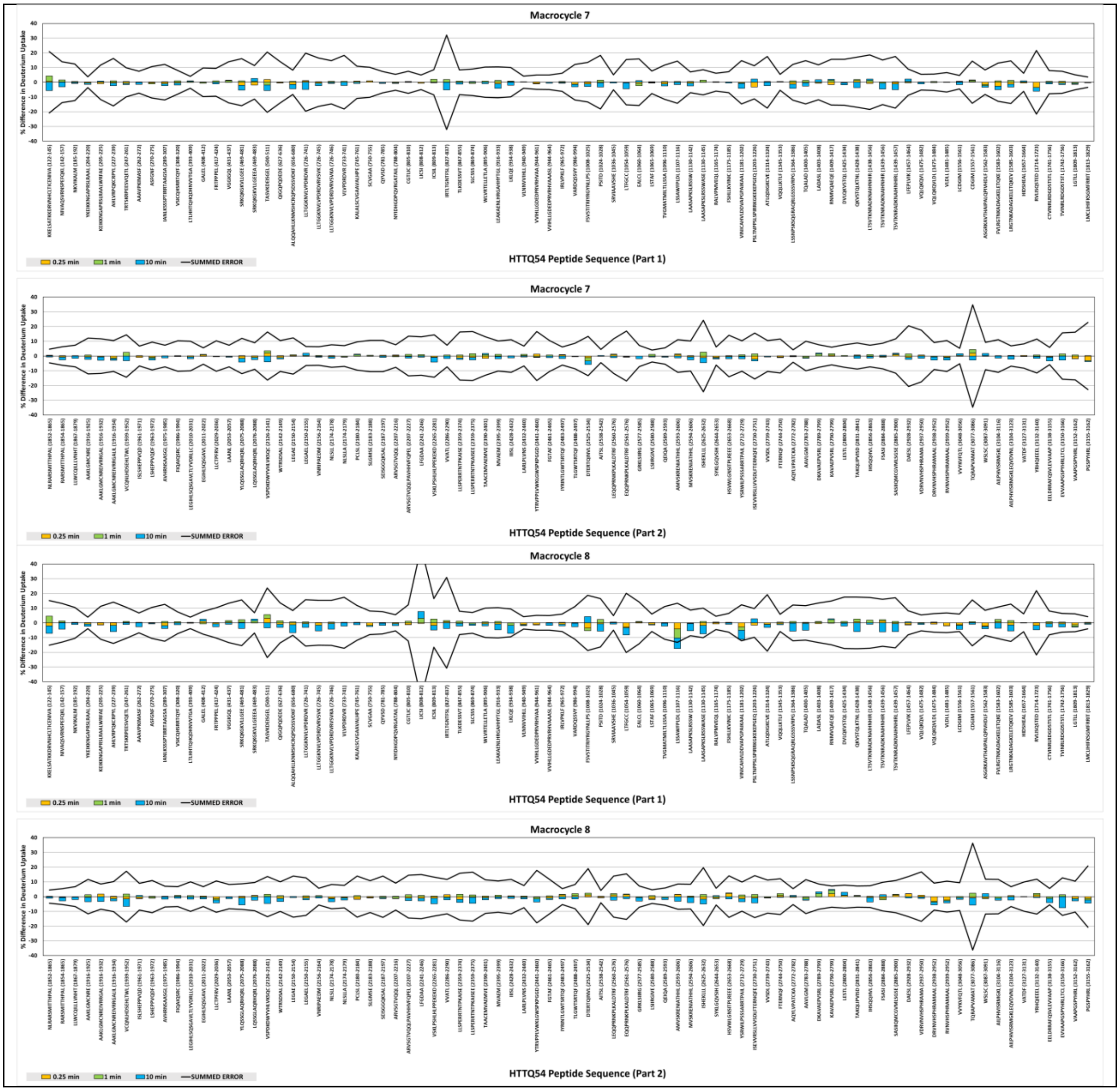


**Supplementary Figure 10.** HTT Timepoint-Level ΔHDX-MS

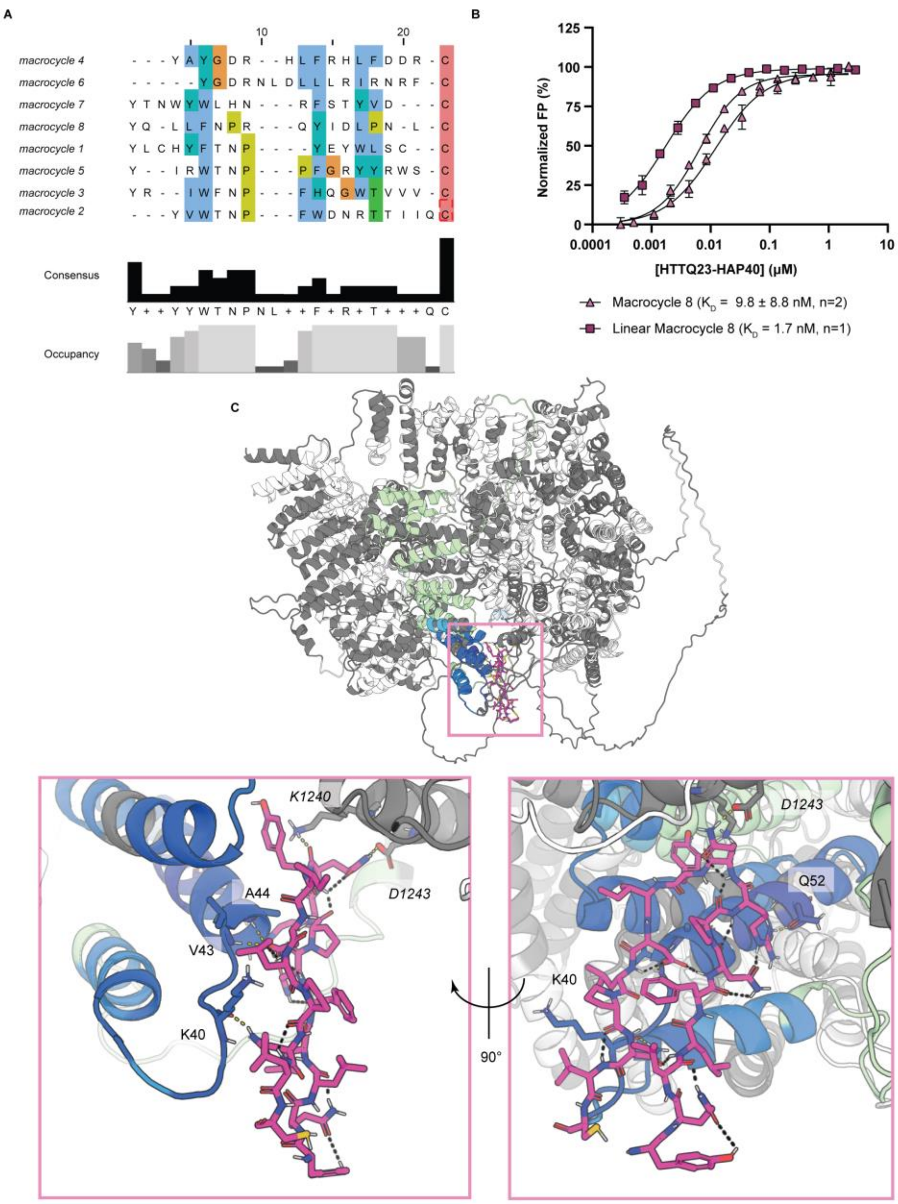


**Supplementary Figure 11.** HADDOCK modeled HTTQ23-HAP40 + linear **8**. **A**, Sequence aligned macrocycles (ClustalX).[42] **B**, FP of FITC-labeled linear **8** compared to cyclic **8**. **C**, HADDOCK model overlayed with HTT-HAP40 HDX heatmap of macrocycle **8**. Intramolecular and intermolecular bonds are shown as black and yellow dashed lines, respectively, while HAP40, HTT, and **8** are green, white (italicized), and pink, respectively. Regions with decreased deuterium uptake are blue while those which did not have sequence coverage in HDX-MS are coloured grey.

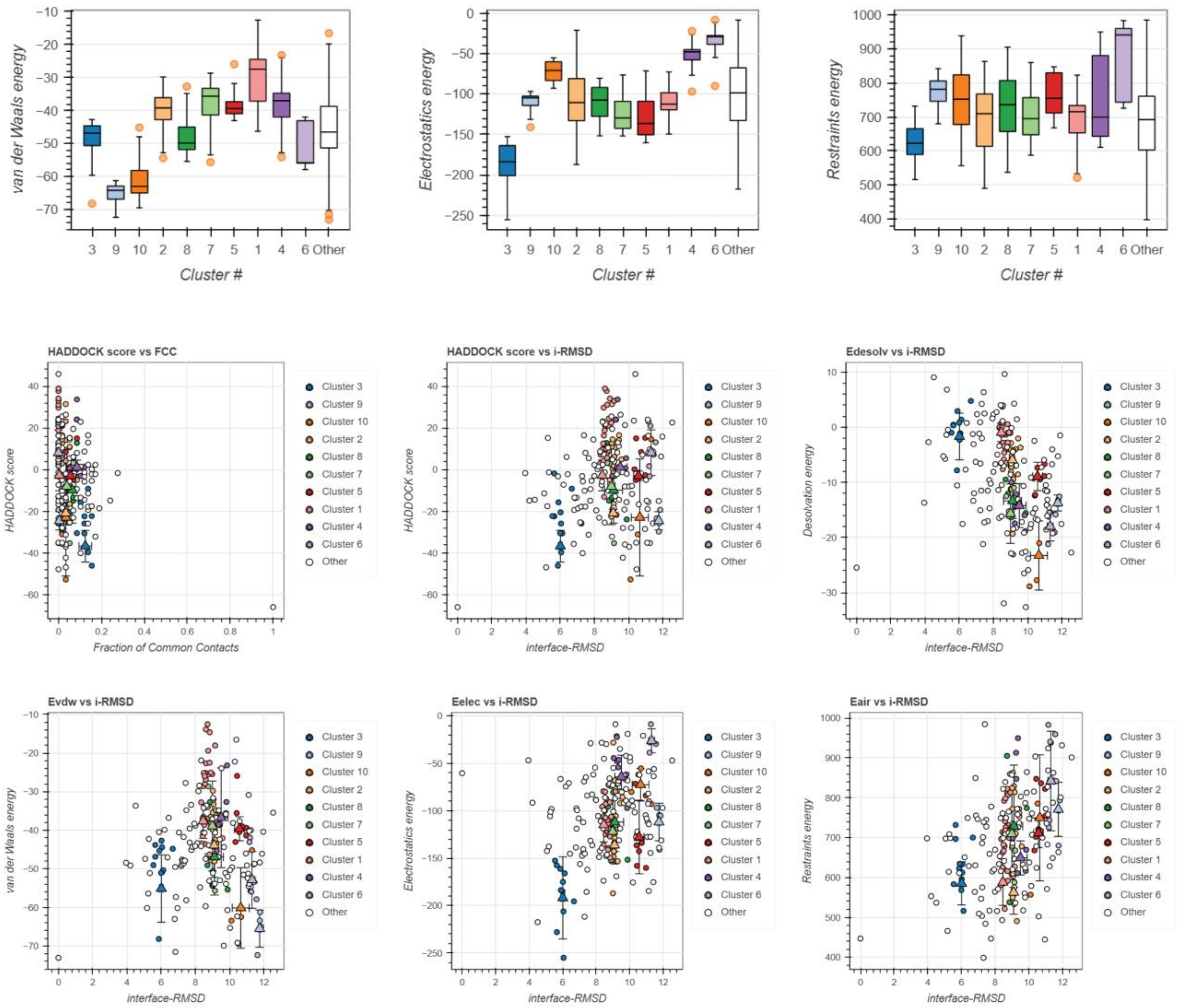

**Supplementary Figure 12.** HADDOCK Cluster Scoring. FCC – fraction of common contacts, i-RMSD – interface RMSD, Edesolv – Desolvation energy, Evdw – intermolecular van der Waals energy, Eelec - intermolecular electrostatic energy, Eair – Ambiguous Interaction Restraints Energy.

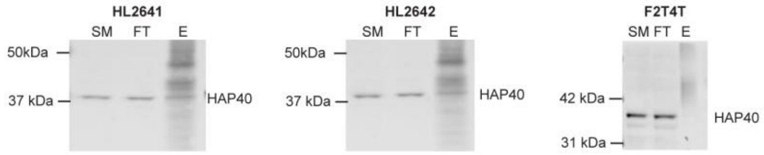


**Supplementary Figure 13. Immunoprecipitation-western blot (IP-WB) analysis of commercially available HAP40 monoclonal antibodies.** HAP40 is readily detected in input and flow-through fractions; little to no enrichment is observed in the elution across all antibodies tested. Samples corresponding to starting material (SM) 2%, flow-through (FT) 2%, and elution (E) 100% fractions were analyzed using the anti-HAP40 antibody CH03722.

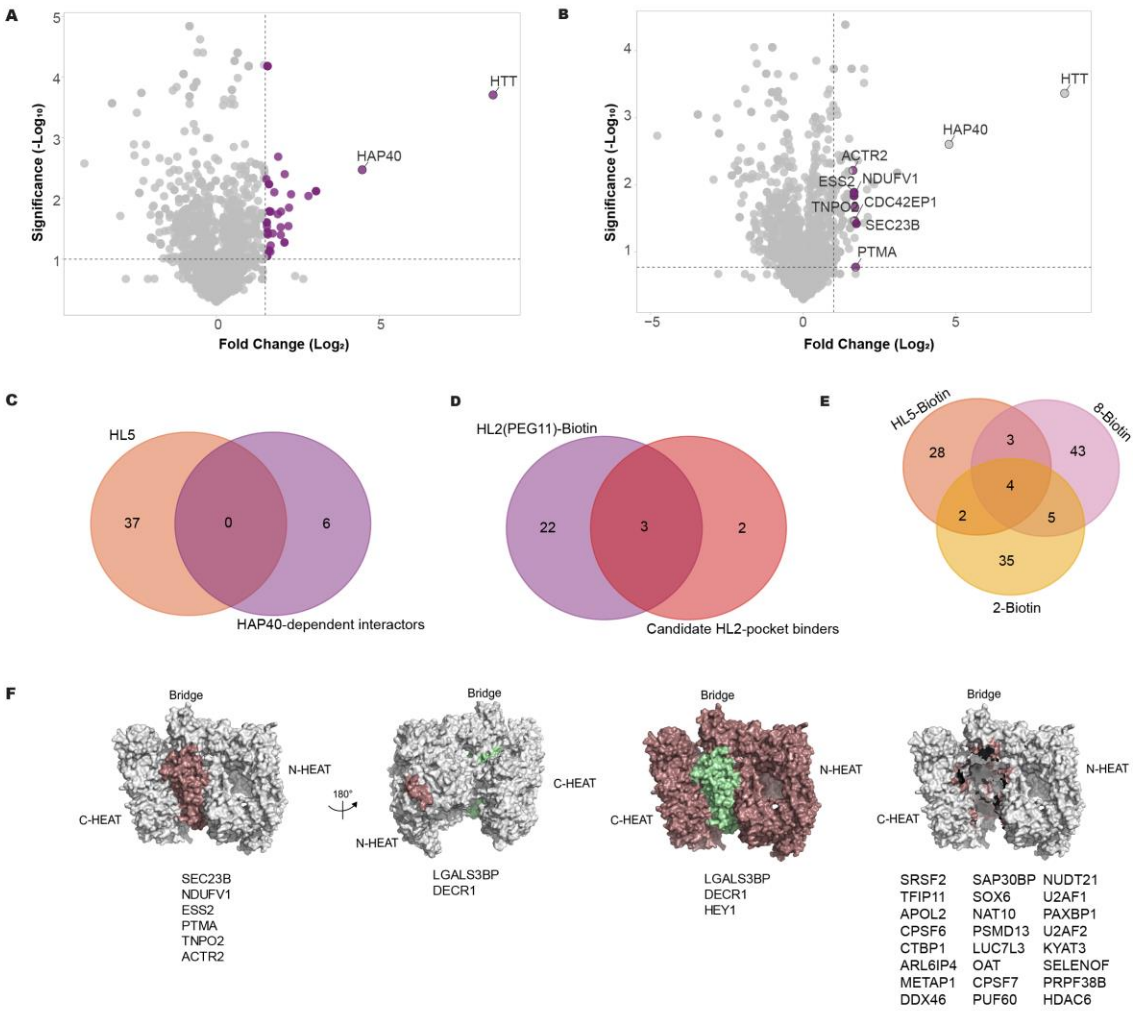


**Supplementary Figure 14. Comparative analysis of macrocycle pulldown datasets.** (A) Volcano plot showing proteins enriched in HL2(PEG11)-Biotin pulldowns in the presence of macrocycle 8. Proteins with ≥3-fold enrichment cutoff relative to HTT knockout are highlighted. (B) Volcano plot of proteins enriched in HL2(PEG11)-Biotin pulldowns in the absence of macrocycle 8. Proteins significantly reduced upon addition of macrocycle 8 are highlighted in purple (**Supplementary Datasheet 2**). The $\log_2$ fold change (WT vs HTT KO) versus $-\log_{10}$ P values derived from normalized protein spectral counts (n= 3). (C) Venn diagram showing overlap between proteins enriched using the apo HTT-binding macrocycle HL5-Biotin and those identified as HAP40-dependent interactors. (D) Venn diagram comparing proteins enriched in HL2(PEG11)-Biotin pulldowns with candidate proteins associated with the HTT N-terminal region. (E) Overlap of proteins identified across pulldowns using HAP40-targeting macrocycles (2-Biotin and 8-Biotin) and the HTT-targeting macrocycle HL5-Biotin. (F) Structural mapping of representative protein subsets onto the HTT–HAP40 complex. Far Left: proteins selectively lost upon macrocycle 8 treatment and not enriched across conditions, consistent with HAP40-dependent associations. Middle left: proteins enriched by HAP40-targeting macrocycles but not overlapping with HL2 datasets, suggesting binding proximal to the N-terminal region. Middle Right: Proteins potentially interacting with diverse surfaces of the HTT-HAP40 complex Far right: Proteins uniquely enriched by HL5 (apo HTT binder) are shown, suggesting preferential interaction with regions inaccessible in the HAP40-bound state.
